# Age-related sensory dysfunction reconfigures the spinal circuitry for touch, itch and pain

**DOI:** 10.64898/2025.12.18.695289

**Authors:** David Acton, Sofia Pimpinella, Martyn Goulding

## Abstract

Sensory dysfunction during aging results in sharp increases in the incidence of chronic itch and pain, which counterintuitively are associated with reduced sensitivity to light touch. While age-related changes in peripheral mechanosensory transmission have been described, the contribution central mechanisms make to altered itch and pain responses remains largely unknown. Here, we show that signalling from cutaneous touch receptors is essential for maintaining the cellular composition of the central somatosensory circuits that process itch and pain information in adult mice. In particular, the loss of excitatory signalling from Merkel cells during aging causes the degeneration of subsets of molecularly defined inhibitory and excitatory neurons in the dorsal spinal cord, including inhibitory neurons that express the neuropeptide NPY. Our demonstration that the activity-dependent loss of inhibitory NPY neurons drives itch and pain hypersensitivity in old animals reveals the mechanism by which aging reconfigures the central neuronal circuits that sense touch, itch and pain.

## Introduction

Mechanosensory information is transmitted and gated within the dorsal horn of the spinal cord by dedicated circuits composed of functionally and molecularly differentiated populations of interneurons^1,2^. These circuits are specified and wired up prenatally^3,4^ and then refined during the first postnatal weeks. During this critical period, the thresholds for noxious and innocuous touch are adjusted according to the changing needs of the animal in an experience-dependent manner ^5–7^, with light touch information modifying paw withdrawal responses and synaptic connectivity in the spinal reflex pathways of rodents^7,8^ and human infants^9,10^.

The thresholds for touch, itch, pain and thermal perception also undergo changes as animals age. These changes occur concurrently with reduced touch signalling caused by the degeneration of cutaneous mechanosensory end organs^11–14^ and demyelination of the sensory nerves that innervate them ^15^. What is not known is whether and how these age-related changes in peripheral touch transmission alter the central circuits that process somatosensory information, and whether these alterations contribute to age-related itch and pain conditions^16–21^

Here, we show that the survival of neurons in the dorsal horn of adult mice is dependent on the activity of cutaneous low-threshold mechanosensory (LTMR) afferents. We demonstrate that a chronic reduction of input from Merkel cells, as occurs during aging, results in the apoptosis of interneurons in the low-threshold mechanoreceptor recipient zone (LTMR-RZ). Moreover, subsets of molecularly defined populations of dorsal horn neurons are differentially sensitive to the age-related loss of input from Merkel cells, with interneurons expressing neuropeptide Y (NPY^+^ INs) being disproportionately affected. In healthy young adult mice, these NPY^+^ INs gate the transmission of mechanical itch information^22–24^ as well as neuropathic and inflammatory pain^25–27^. We find the loss of NPY^+^ INs that occurs following the reduction of input from Merkel cells during aging results in reduced NPY tone within the dorsal horn and disinhibition of the pathways that transmit mechanical itch and inflammatory pain. Together, our findings describe a mechanism of somatosensory dysfunction in which select molecularly defined populations of dorsal horn neurons are reduced in size following the loss of excitatory drive from touch receptors. The consequent restructuring of somatosensory circuits results in substantial changes to the transmission of somatosensory information within the central nervous system.

## Results

### Acute silencing of Merkel cells increases mechanical thresholds but does not change itch sensitivity

Previous studies have led to the hypothesis that Merkel cells activate the inhibitory spinal neurons that gate central itch transmission pathways in young adult mice^14,28^. In support of this, itch responses are elevated following the spontaneous degeneration of Merkel cells that occurs during aging (Fig. 1a, b)^14^. We set out to further test the contribution that Merkel cells make to the gating of itch by asking if responses to mechanosensory stimuli in young adult mice are altered following the inactivation of Merkel cells^29^. Selective deletion of *Piezo2* from Merkel cells following the administration of tamoxifen to P56 *Atoh1^CreER^; Piezo2^f/f^* mice^29^ reduced their ability to detect indentation of the glabrous skin by the finest von Frey hairs at P70 with no effect on dynamic touch or acute pain responses (Extended Data Fig. 1, a-c). This accords with the previously reported role of Merkel cells in the detection of light static but not dynamic touch^29^. Strikingly, these mice exhibited increased sensitivity to mechanical itch stimulation of the hairy skin of the nape by low-grade von Frey filaments^14^ and an increase in spontaneous scratching, which is consistent with reduced gating of mechanical itch by inhibitory neurons in the spinal cord (Fig 1, C-D).

**Figure 1.**
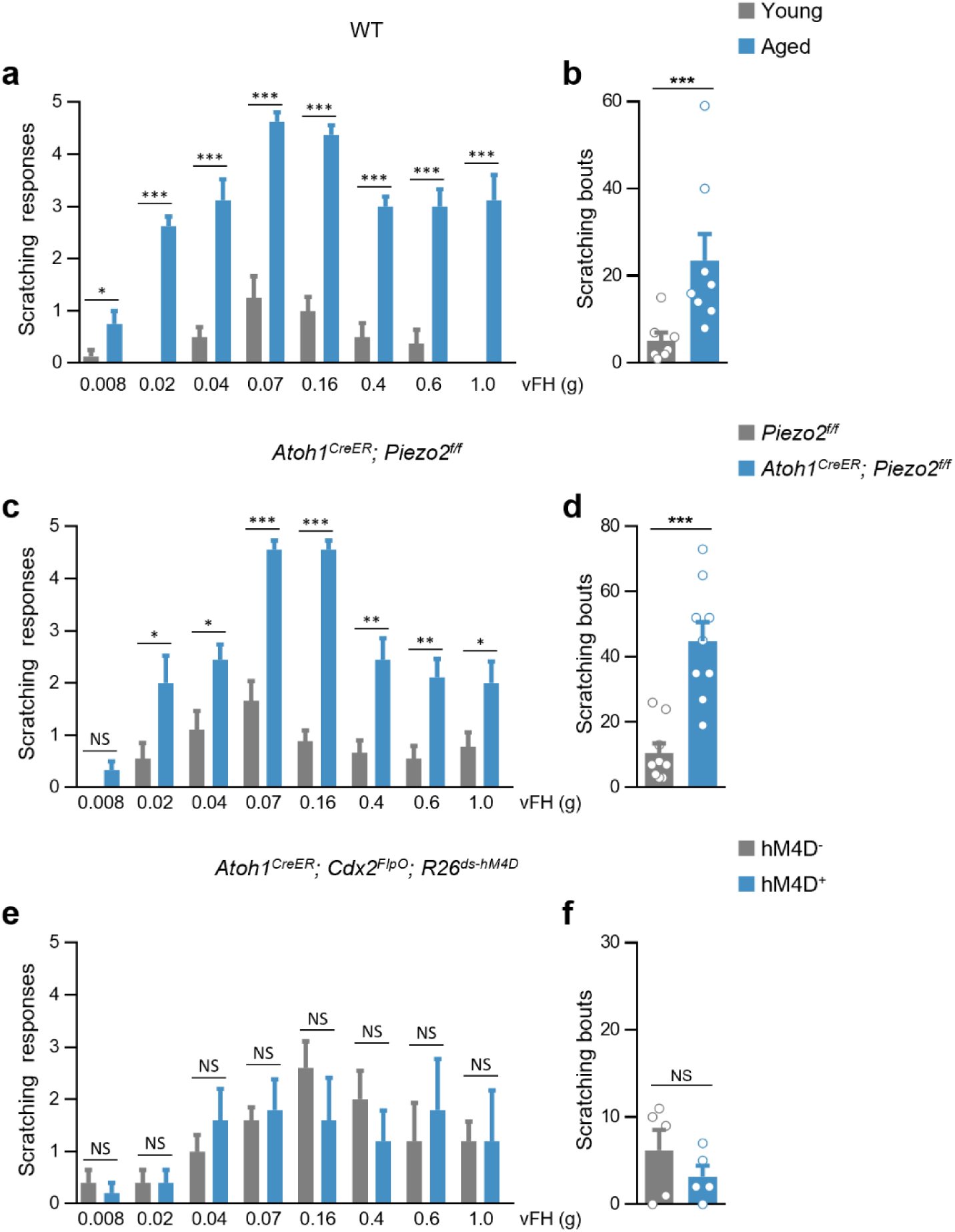
Chronic but not acute loss of Merkel cell function exacerbates itch. **a, b,** Aged (2-year-old) wild type mice (n = 8) exhibit greater responsiveness to mechanical itch stimulation by von Frey hairs applied to the nape (**a**) and greater rates of spontaneous hindlimb scratching recorded over a 30-min period (**b**) than young adult (P56) mice (n = 8). **c, d,** Chronic silencing of Merkel cells at P56 increases mechanical itch sensitivity (**c**) and spontaneous scratching in *Atoh1^CreER^; Piezo2^f/f^* mice (n = 9) as compared to control Piezo2^f^*^/f^* mice (n = 9). **e, f,** Acute chemogenetic silencing of Merkel cells in CNO-treated *Atoh1^CreER^; Cdx2^FlpO^; R26^ds-hM4D^* mice (n =5) has no effect on mechanical (**e**) or spontaneous itch (**f**) responses as compared to CNO-treated *Atoh1^CreER^; R26^ds-hM4D^* controls (n =5).

Conflicting reports regarding the normal function of Merkel cells in sensory discrimination suggest that plasticity within central circuits might mask their normal function following chronic loss-of-function manipulations^29–31^. To address this possibility, we tested itch responses in mice following the acute silencing of Merkel cells. In *Atoh1^CreER^; Cdx2::FlpO; R26^ds-hM4D^* mice treated with tamoxifen at P56, expression of the inhibitory DREADD hM4D is restricted to Merkel cells in the skin (Extended Data Fig. 1d, e), and is undetectable in the DRG, spinal cord and brain (n = 5 mice). Acute silencing of Merkel cells following injection of CNO at P70 substantially reduced the sensitivity of the glabrous skin to punctate touch stimulation without affecting dynamic touch or acute pain responses, similar to what occurs after the chronic silencing of Merkel cells (Extended Data Fig. 1, f-h). However, acute silencing of Merkel cells in *Atoh1^CreER^; Cdx2::FlpO; R26^ds-hM4D^* mice had no effect on either mechanical itch sensitivity or spontaneous scratching as compared to control *Atoh1^CreER^; R26^ds-hM4D^* mice that were treated identically (Fig. 1e, f). Together these data show that the chronic but not the acute loss of Merkel cell activity gives rise to hyperknesis, indicating that Merkel cells alone do not drive the inhibitory neurons in the spinal cord that gate mechanical itch. It therefore appears that the changes in itch sensitivity observed following the persistent loss of input from Merkel cells, both in aged mice and following the chronic deletion of *Piezo2*, are instead driven by long-term changes to the central circuits that gate mechanical itch.

### Activity-dependent survival of dorsal horn neurons in adult mice

As an initial step towards assessing how the persistent loss of input from Merkel cells alters the spinal circuits that transmit itch, we analysed the dorsal horn of mice at 2 years of age, a timepoint when Merkel cells have degenerated and activity in the slowly adapting type 1 (SAI) fibres that innervate them is severely depressed^14^. Aged mice displayed a large reduction (23.7 ± 3.7%) in the total number of Neurotrace^+^ neurons compared to P56 young adult mice (p < 0.01; aged, n = 5 mice; young, n = 4 mice). Notably, this loss of neurons was confined to the low-threshold mechanoreceptor recipient zone (LTMR-RZ; laminae IIi-V), with the greatest reduction occurring in laminae IIi and III, (Fig. 2a, b). The loss of LTMR-RZ neurons in aged mice was caused by a reduction in both *Slc32a1* (VGAT)-expressing inhibitory neurons (-30.3 ± 6.1%) and *Slc17α6* (vGluT2)-expressing excitatory neurons (-33.2 ± 2.4%; Fig. 2c, d). Cells displaying TUNEL-reactive double-strand DNA breaks and activated caspase-3 immunolabeling, while rare in sections from the dorsal horn of young mice, were frequently observed in the dorsal horn of aged mice (Extended Data Fig. 2, a-c; Fig. 2e, f). Together, these findings argue that neurons in the LTMR-RZ undergo programmed cell death as animals age and suggest that neuronal cell loss may underlie the dysfunctional processing of somatosensory information in older animals.

**Figure 2:**
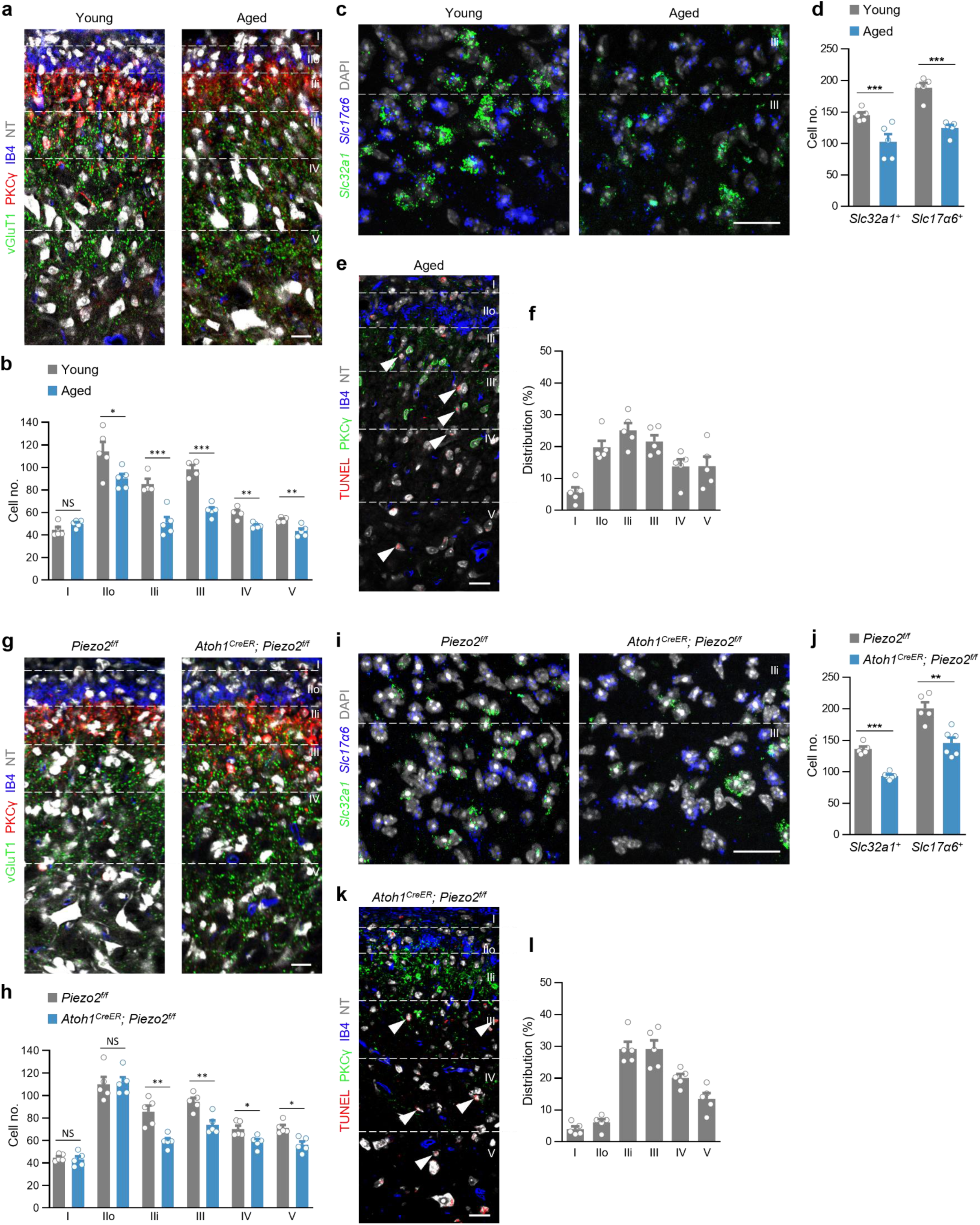
Activity-dependent and age-related cell survival in the dorsal horn. **a,** Sections through the dorsal horn at C5 of young (left) and aged mice showing NeuroTrace (NT)-stained neurons; vGluT1, PKCγ, and IB4 counterstaining are also shown to indicate laminar boundaries. **b,** Reduced neuron number in the LTMR-RZ (lamina IIi-V) of aged compared to young mice. **c, d,** Representative dorsal horn sections (**c**) and summary plot (**d**) showing a reduction in the number of inhibitory neurons expressing *Slc32a1* mRNA and excitatory neurons expressing *Slc17α6* in aged compared to young mice. **e, f,** Representative dorsal horn section showing TUNEL reactivity in neurons (arrowheads; **e**) and summary of the laminar distribution of TUNEL^+^ neurons (**f**) in aged mice. **g,** Sections through the dorsal horn at C5 of control (left) and Merkel cell-silenced mice showing NT-stained neurons. (H) Reduced neuron number in the LTMR-RZ (lamina IIi-V) of Merkel cell-silenced mice compared to controls. **i, j,** Representative dorsal horn sections (**i**) and summary plot (**j**) showing a reduction in the number of inhibitory *Slc32a1^+^* and excitatory *Slc17α6^+^* neurons following Merkel cell silencing. **k, l,** Dorsal horn section showing TUNEL reactivity in neurons (arrowheads; **k**) and summary of the distribution of TUNEL^+^ neurons (**l**) following Merkel cell silencing. Scale bars, 20 µm. Error bars represent SEM. *p < 0.05; **p < 0.01; ***p < 0.001; ns, no significant difference.

We then tested the hypothesis that this cell loss is due to reduced input from Merkel cells by analysing the dorsal horn of young adult mice in which Merkel cells were chronically inactivated. *Atoh1^CreER^; Piezo2^f/f^* mice treated with tamoxifen at P56 displayed a marked reduction in Neurotrace^+^ neurons (20.3 ± 3.7%) in the dorsal horn compared to tamoxifen-treated *Piezo2^f/f^* control littermates (p < 0.001; silenced versus control mice, n = 5 mice in each cohort). This loss of neurons was confined to the LTMR-RZ, where SAI fibres terminate (Brown, 1981; Abraira and Ginty, 2013), with the greatest losses occurring in laminae IIi and III. By contrast, there was little or no loss of neurons in the superficial laminae (laminae I-IIo) innervated by nociceptive afferents (Fig. 2g, h). This preferential loss of neurons in the LTMR-RZ involved both *Slc32a1^+^* inhibitory (- 31.4 ± 1.6%) and *Slc17α6^+^* excitatory neurons (-27.3 ± 4.2%; Fig. 2i, j), phenocopying the neuronal loss that is seen in aged mice. Consistent with these data, an increase in TUNEL^+^ and activated Caspase-3-reactive nuclei was observed in dorsal horn sections from Merkel cell-silenced mice but not control mice (Extended Data Fig. 2e, f). As expected, the vast majority of TUNEL^+^ neurons were located in laminae IIi-V, the region of the LTMR-RZ that is innervated by Merkel cell-SAI afferents (Fig. 2k, l). Taken together, these findings establish that ongoing Merkel cell activity supports the viability of dorsal horn neurons in adult mice.

### Molecularly defined populations of inhibitory neurons are differentially susceptible to apoptosis following reduced input from Merkel cells

We next sought to understand how the activity-dependent and age-related loss of neurons in the dorsal horn affects the processing of somatosensory information. In view of our finding that the gating of itch transmission at the spinal level is impaired following the chronic loss of input from Merkel cells, we performed a detailed analysis of the effects of the loss of Merkel cell input on molecularly defined populations of inhibitory spinal interneurons, focusing on those associated with the gating of itch.

Pax2 selectively marks all inhibitory neurons in the dorsal horn, including those subpopulations that gate itch and pain transmission^1,3,32,33^. Cell counts in 2-year-old mice revealed a marked reduction in Pax2-expressing neurons (29.0 ± 3.7%) compared to young mice. This was accompanied by a pronounced increase in Pax2^+^/TUNEL^+^ cells (Fig. 3, a-c), with Pax2^+^/TUNEL^+^ neurons representing 41.4 ± 4.5% of all TUNEL-labelled neurons in the dorsal horn. A comparable reduction in immunolabeled Pax2^+^ neurons (28.6 ± 3.7%) was seen in *Atoh1^CreER^; Piezo2^f/f^* mice, with TUNEL labelling present in a significant fraction of the remaining Pax2^+^ cells (Fig. 3, d-f). Importantly, 43.5 ± 2.2% TUNEL^+^ neurons showed Pax2 immunoreactivity, providing further evidence that the survival of inhibitory dorsal horn neurons requires ongoing excitatory drive from Merkel cells.

**Figure 3:**
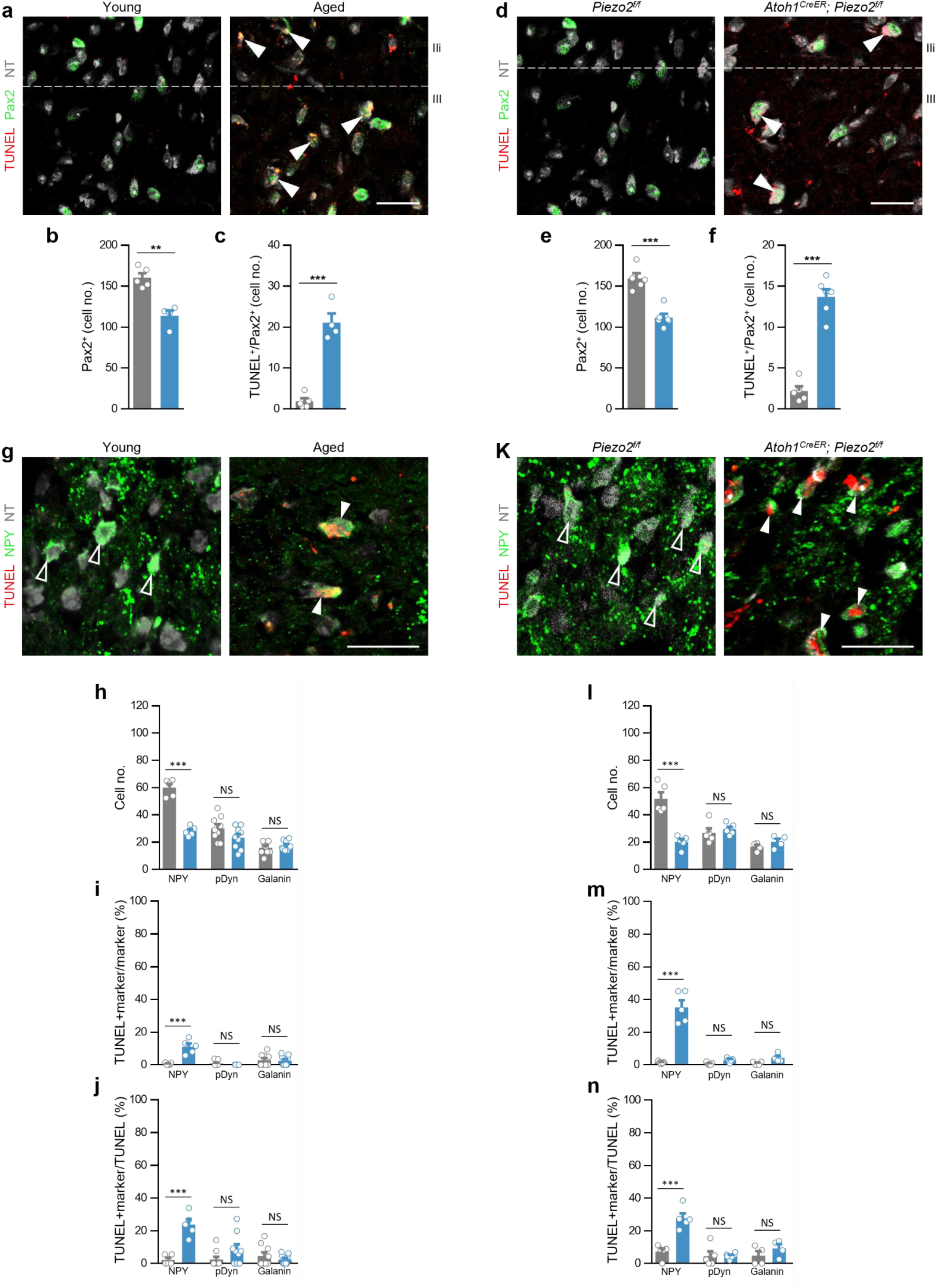
Differential loss of Pax2-derived neurons following loss of Merkel cell activity. **a,** Example images showing TUNEL labelling in Pax2^+^ neurons (arrowheads) in young (left) and aged mice (right). **b, c,** Summary plots showing a reduction in Pax2^+^ neurons per section (**b**) and a corresponding increase in TUNEL^+^/Pax2^+^ neurons per section in aged mice (**c**). **d-f,** TUNEL labelling (**d**) in Pax2^+^ neurons (arrowheads) in control (left) and Merkel cell-silenced mice (right) and summary plots showing a reduction in Pax2^+^ neurons per section in Merkel cell-silenced mice (**e**) and a corresponding increase in TUNEL^+^/Pax2^+^ neurons per section (**f**). **g,** Representative images showing TUNEL staining in NPY-immunoreactive, NeuroTrace (NT)-labelled neurons in the dorsal horn of aged (right) but not young mice (left). **h-j,** Assessment of apoptotic cell loss in populations of molecularly defined inhibitory neurons identified by antibodies against NPY, dynorphin (prodynorphin; pDyn), and galanin in aged compared to young mice: summaries of population size (**h**), the proportion of TUNEL^+^ neurons expressing each molecular marker (**i**), and the proportion of each molecularly defined population labelled by TUNEL (**j**). **k,** Representative images showing TUNEL staining in NPY-immunoreactive, NT-labelled neurons in the dorsal horn of Merkel cell-silenced (right) but not control mice (left). **l-n,** Programmed cell death in populations of molecularly defined inhibitory neurons, identified by immunohistochemistry, in Merkel cell-silenced mice compared to controls: summaries of population size (**l**), the proportion of TUNEL^+^ neurons expressing each molecular marker (**m**), and the proportion of each molecularly defined population labelled by TUNEL (**n**). Scale bars, 20 µm. Error bars represent SEM. *p < 0.05; **p < 0.01; ***p < 0.001; ns, no significant difference.

Two prominent subpopulations of inhibitory neurons specified by Pax2 have been shown to modulate itch^23,34–37^. These are the dynorphin^+^ neurons that gate chemically evoked and morphine-induced itch^37–39^ and the NPY^+^ INs that selectively gate mechanical itch, as well as some forms of pain^22–24,40^. No reduction was seen in the dynorphin^+^ population either in aged mice compared to young mice or following the experimental silencing of Merkel cells (Fig. 3, g-n; Extended Data Fig 3a, b, h, i). Likewise, there was no change in the galanin-expressing population with which it overlaps. Moreover, very few of the dynorphin^+^ and galanin^+^ neurons exhibited TUNEL-labelling after removing Merkel cells.

By contrast, we observed marked reductions in the number of interneurons expressing NPY in the dorsal horn of mice lacking input from Merkel cells, with NPY-immunoreactive neurons reduced by 53.3 ± 2.2% in aged mice compared to young controls and by 60.8 ± 3.3% in *Atoh1^CreER^; Piezo2^f/f^* mice compared to controls (Fig. 3, g-n). Interestingly, this loss of NPY^+^ INs is comparable to the 60-65% reduction in NPY^+^ INs that is seen following diphtheria toxin receptor-mediated ablation, which also causes mechanical itch^22,23^. This loss of NPY^+^ INs also accounts for the reduction in NPY immunofluorescence and reduced expression of *Npy* mRNA that occurs in the dorsal horn of aged mice (Ref. 41; this study) and in Merkel cell-silenced mice (Extended Data Fig. 3, c-f, j-m). Our observation that a large fraction of TUNEL^+^ neurons in the dorsal horn of aged and Merkel cell-silenced mice expresses NPY (Fig. 3, g-n) but not dynorphin or galanin demonstrates that functionally specialized populations of dorsal horn inhibitory interneurons are differentially sensitive to reduced input from Merkel cells. Moreover, our discovery that the survival of NPY^+^ INs neurons requires excitatory input from Merkel cells, coupled with monosynaptic rabies tracing experiments^23,42–44^ showing NPY^+^/Lbx1^+^ neurons are innervated by NF200^+^ fibres closely opposed to K8^+^ Merkel cells throughout the hairy skin of the nape (n = 5 mice; Extended Data Fig. 4), provides evidence for the direct innervation of NPY^+^ INs by Merkel cell-SAI neurite complexes.

### Activity-dependent degeneration of NPY^+^ INs exacerbates itch during normal aging

To address the functional consequences of the activity-dependent cell loss that occurs during aging, we asked whether the loss of NPY^+^ INs underlies the increases in spontaneous and mechanically evoked itch observed following the degeneration of Merkel cells in aged mice. The NPY^+^ INs gate the transmission pathway for mechanical itch, which is mediated by Calcrl^+^/Gpr83^+^ neurons that project to the parabrachial nucleus^22–24^. Consequently, we hypothesized that selectively ablating the Calcrl^+^/Gpr83^+^ projection neurons would alleviate the increased sensitivity to mechanical itch seen in aged mice. As predicted, ablation of the Calcrl^+^/Gpr83^+^ neurons using PEN-SAP^24^ reversed both the mechanical itch hypersensitivity and elevated levels of spontaneous scratching observed in aged control mice (Fig. 4a, b), whereas ablating the spinal Tacr1^+^ projection neurons that transmit acute chemical itch had no effect (Fig. 4c, d)^22,24,45^.

**Figure 4:**
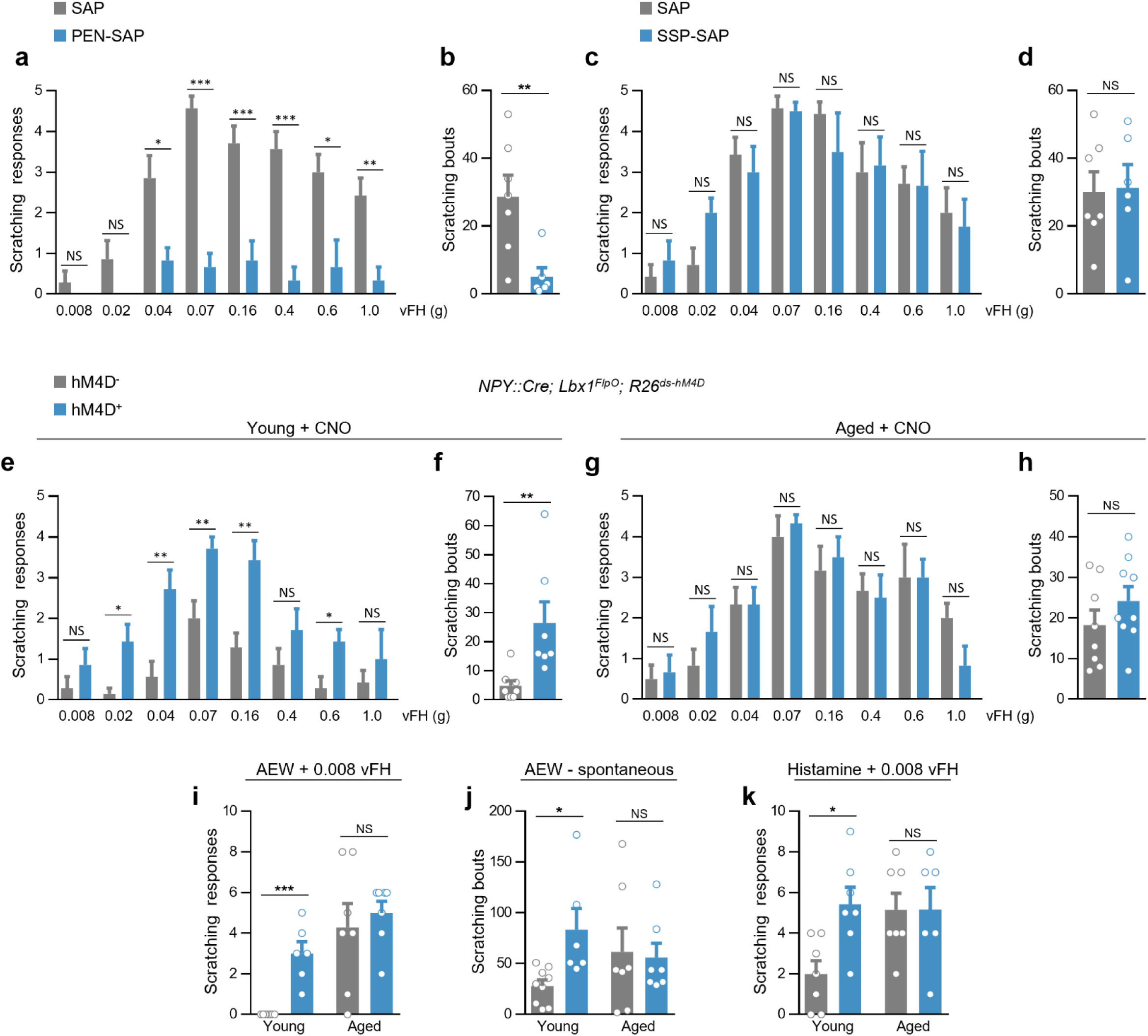
NPY^+^ INs do not gate itch following Merkel cell degeneration in aged mice. **a, b,** In aged mice, ablation of Gpr83^+^ neurons by injection of PEN-SAP reverses mechanical hyperknesis (**a**; control blank SAP, n = 7; PEN-SAP, n =6) and prevents spontaneous scratching (**b**). **c, d,** Ablation of Tacr1^+^ neurons in aged mice by injection of SSP-SAP has no effect on mechanical itch sensitivity (**c**; SAP, n = 7; SSP-SAP, n =6) or spontaneous scratching (**d**). **e, f,** Mechanical itch sensitivity (**e**) and spontaneous scratching (**f**) are increased by administration of CNO to P56 *NPY::Cre; Lbx1^FlpO^; R26^ds-hM4D^* mice (hM4D^+^; n = 7) as compared to control *NPY::Cre; R26^ds-hM4D^* mice (hM4D^-^; n = 7). **g, h,** Mechanical itch sensitivity (**g**) and spontaneous scratching (**h**) are unchanged following administration of CNO to aged *NPY::Cre; Lbx1^FlpO^; R26^ds-hM4D^* mice (hM4D^+^; n = 6) compared to aged control *NPY::Cre; R26^ds-hM4D^* mice (hM4D^-^; n = 6). **i, j,** Chemogenetic silencing of spinal NPY^+^ neurons potentiates hyperknesis/alloknesis (**i**) and spontaneous scratching (**j**) in young but not aged mice treated with AEW to induce dry skin itch. **k,** Silencing spinal NPY^+^ neurons exacerbates histamine alloknesis in young but not aged mice. Error bars represent SEM. *p < 0.05; **p < 0.01; ***p < 0.001; ns, no significant difference.

We then tested a second prediction, namely that silencing the NPY^+^ INs in aged animals would have a smaller effect on acute itch responses than in young animals, reflecting a smaller contribution of the NPY^+^ IN population to the gating of itch in aged mice. As previously reported^23^, chemogenetic silencing of the NPY^+^ INs in young *NPY::Cre; Lbx1^FlpO^; R26^ds-hM4D^* mice caused a marked sensitization to mechanical itch, as well as increased levels of spontaneous scratching (Fig. 4e, f). By contrast, silencing NPY^+^ INs in aged mice had no detectable effect on either mechanical or spontaneous itch (Fig. 4g, h), which is consistent with the loss of inhibition of mechanical itch by NPY^+^ INs in aged mice.

The mechanical itch transmission pathway gated by NPY^+^ INs also contributes to pathological itch^24,40^, prompting us to assess whether chronic itch and acute hyperkinesis are altered between young and aged mice. In the AEW model of chronic dry skin itch^46,47^, silencing NPY^+^ INs in young mice exacerbated both hyperknesis and spontaneous scratching but had no effect on either measure in aged mice (Fig. 4i, j). Similarly, young but not aged mice were sensitized to tactile stimuli following the silencing of NPY^+^ INs in a modified model of acute hyperknesis. In this model, intradermal (i.d.) injection of the nape with histamine sensitizes the surrounding skin to punctate mechanical stimuli^48^. Whereas a 0.008 g filament seldom elicited scratching in young mice, it did so in approximately half of trials in aged mice, demonstrating an increase in hyperknesis. Silencing the NPY^+^ INs in young mice led to scratching at a rate comparable with aged mice, whereas silencing them in aged mice had no effect on the rate of scratching (Fig. 4k). These data indicate that NPY^+^ INs modulate the mechanical itch transmission pathway in young adult mice but not in geriatric mice, which is consistent with the reduction in NPY^+^ INs seen in aged mice. Taken together these findings demonstrate that the central somatosensory circuitry for itch is reconfigured in aged mice.

### NPY-Y1 signalling gates itch in young adult but not aged mice

Our prior analysis has shown that mechanical itch is principally gated by NPY signalling via the neuropeptide Y receptor type 1 (Y1) as opposed to fast synaptic inhibition^22^, thus raising the possibility that the increase in mechanical itch sensitivity during aging is due to the loss of inhibitory NPY-Y1 signalling. To test this, we first assessed the effects of silencing NPY^+^ INs while blocking Y1 activity. When young adult *NPY::Cre; Lbx1^FlpO^; R26^ds-hM4D^* mice were pretreated with BIBP 3226 to selectively block Y1 receptors^22,49^, the silencing of NPY^+^ INs with CNO failed to further increase mechanical itch sensitivity and spontaneous scratching (Fig. 5, a-d). These data corroborate our previous finding that in young adult mice the inhibitory actions of the NPY^+^ INs are mediated predominantly by a peptidergic mechanism, being consistent with the effects of selectively inactivating the Y1 receptor in the dorsal spinal cord^22^.

**Figure 5:**
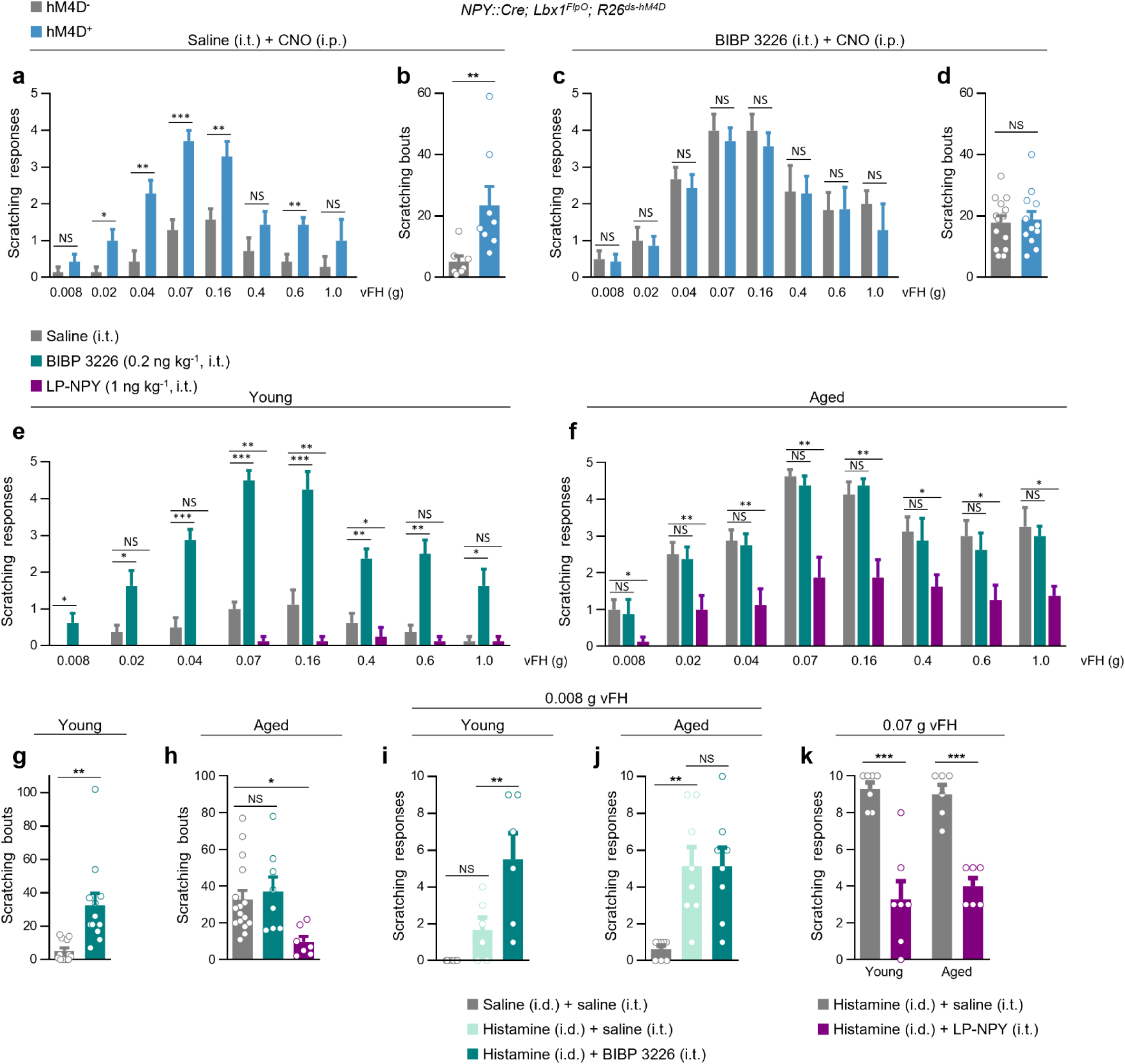
NPY-Y1 signalling does not gate itch in aged mice following Merkel cell degeneration. **a, b,** Mechanical itch responsiveness (**a**; *NPY::Cre; R26^ds-hM4D^* controls, n = 6; *NPY::Cre; Lbx1^FlpO^; R26^ds-hM4D^*, n = 7) and spontaneous scratching (**b**) are elevated following the silencing of spinal NPY^+^ neurons when mice are pretreated with control saline. **c, d,** Pretreatment of mice with the selective Y1 antagonist BIBP 3226 (0.2 ng kg^-1^, i.t.) prevents increases in mechanical itch (**c**; *NPY::Cre; R26^ds-hM4D^* controls, n = 6; *NPY::Cre; Lbx1^FlpO^; R26^ds-hM4D^*, n = 7) and spontaneous scratching (**d**). **e,** Mechanical itch sensitivity in young mice increases following administration of the selective Y1 antagonist BIBP 3226 (0.2 ng kg^-1^, i.t.) compared to control vehicle, but decreases following administration of the selective Y1 agonist [Leu^31^, Pro^34^]-NPY (LP-NPY; 1 ng kg^-1^, i.t.; n = 8). **f,** Mechanical itch sensitivity is not increased by Y1 blockade in aged mice but is suppressed by Y1 activation (n = 8). **g,** Spontaneous scratching is increased by Y1 blockade in young mice. **h,** Spontaneous scratching is not increased by Y1 blockade in aged mice, but it is suppressed by Y1 activation. **i,** Alloknesis, assessed by application of a 0.008 g von Frey hair to the skin around the site of a histamine injection (50 µg in 10 µl, i.d.), is potentiated by Y1 blockade in young mice. **j,** Histamine hyperknesis/alloknesis is not potentiated by Y1 blockade in aged mice. **k,** Y1 activation suppresses histamine hyperknesis/alloknesis, assessed with a 0.07 g von Frey hair in young or aged mice. Error bars represent SEM. *p < 0.05; **p < 0.01; ***p < 0.001; ns, no significant difference.

We then compared the gating of the mechanical itch transmission pathway by NPY-Y1 signalling between young and aged mice. Whereas in young mice pharmacological blockade of Y1 receptors potentiated responses to mechanical itch stimuli and caused spontaneous scratching, in aged mice itch these responses were unchanged (Fig 5. E-H). Similarly, blocking NPY-Y1 signalling with BIBP 3226 exacerbated acute histamine-induced hyperknesis in young mice, such that they now scratched at rates comparable with vehicle-treated aged mice, but aged mice showed no further sensitization after Y1 blockade (Fig. 5i, j). By contrast, activation of Y1 receptors by the selective Y1 agonist [Leu^31^, Pro^34^]-NPY (LP-NPY)^22,50^ suppressed mechanically evoked scratching in both young and aged mice (Fig. 5e, f). Furthermore, it inhibited the spontaneous scratching and histamine hyperknesis seen in aged mice (Fig. 5h, k), arguing that functional Y1 receptors continue to be expressed during aging.

We then tested whether the degeneration of NPY^+^ INs that occurs following the loss of Merkel cell input reduces the gating of itch transmission by NPY-Y1 signalling. BIBP 3226 increased scratching in control but not Merkel cell-silenced mice (Fig. 6, a-d), which supports our model that reduced NPY tone in the dorsal horn of mice following Merkel cell silencing is the primary driver of increased itch sensitivity. Consistent with this, both spontaneous scratching and histamine alloknesis were potentiated in control mice but not in Merkel cell-silenced mice (Fig. 6e, f). In a further test of our model, Y1 receptor activation reduced both touch-evoked and spontaneous scratching in Merkel cell-silenced mice (Fig. 6a, b, d), indicating that the expression of functional Y1 receptors is maintained in these mice. Together, these experiments demonstrate that NPY tone within the dorsal horn is reduced following the loss of input from Merkel cells during aging, with the degeneration of NPY^+^ INs and loss of NPY-Y1 signalling exacerbating itch in multiple acute and pathological contexts.

**Figure 6:**
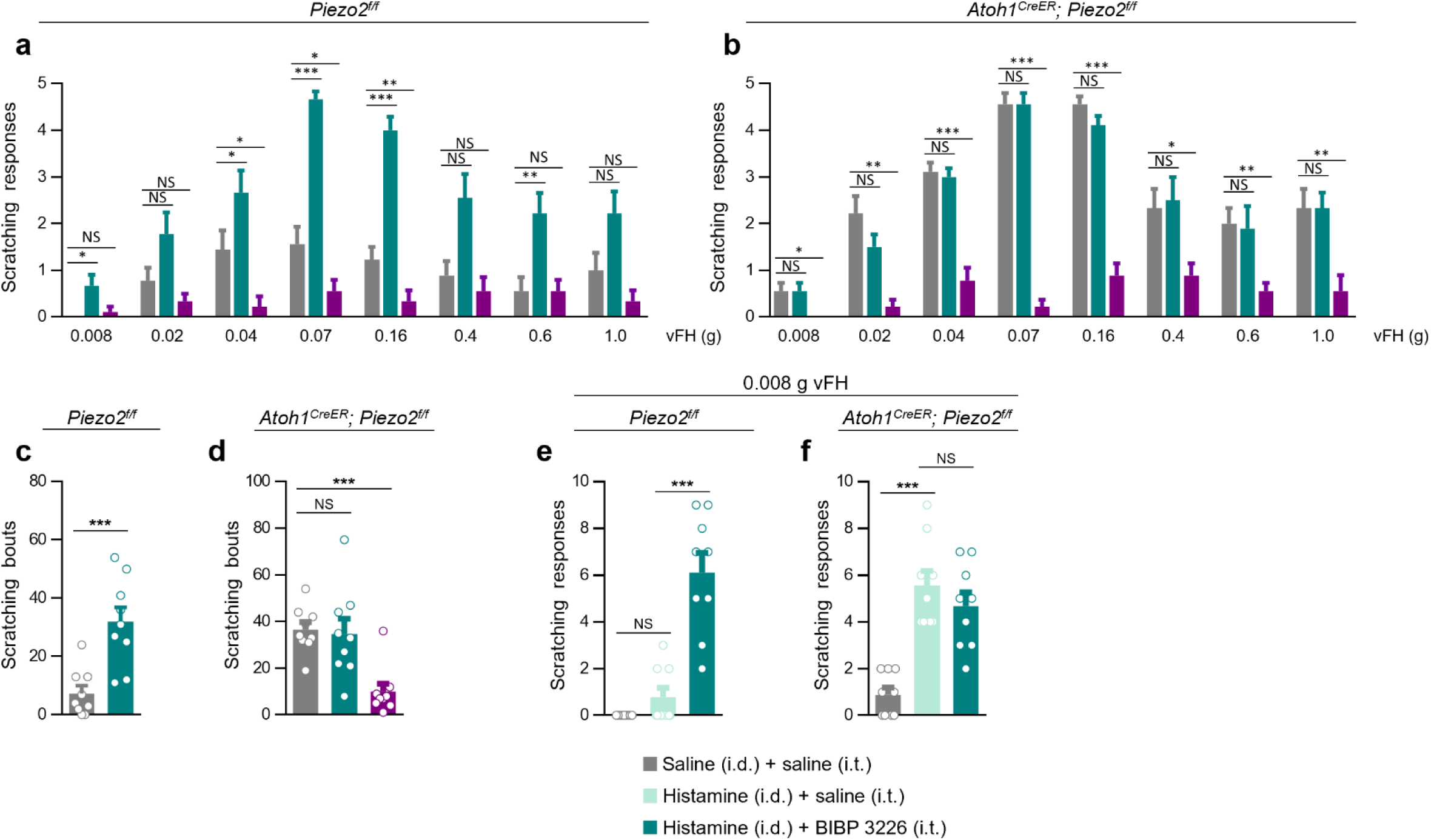
NPY-Y1 signalling does not gate itch following chronic Merkel cell silencing. **a,** Mechanical itch sensitivity increases in *Piezo2^f/f^* control mice following Y1 blockade but decreases following Y1 activation (n = 9). **b,** Mechanical itch sensitivity is not increased by Y1 blockade when Merkel cells are silenced in *Atoh1^CreER^; Piezo2^f/f^* mice, but it is suppressed by Y1 activation (n = 9). **c,** Spontaneous scratching is potentiated by Y1 blockade in control mice. **d,** Spontaneous scratching is unchanged by Y1 blockade when Merkel cells are silenced, but it is suppressed by Y1 activation. **e, f,** Y1 blockade exacerbates histamine hyperknesis/alloknesis in control mice (**e**) but not in mice in which Merkel cells are silenced (**f**). Error bars represent SEM. *p < 0.05; **p < 0.01; ***p < 0.001; ns, no significant difference.

### NPY^+^ INs gate pain in young but not aged mice

In addition to mechanical itch, endogenous NPY-Y1 signalling has been proposed to gate the transmission pathways for inflammatory pain^25,26^. To determine whether the antinociceptive effects of NPY-Y1 signalling reported in young mice are lost during aging following the degeneration of Merkel cells, we assessed thermal nociception in a model of inflammatory pain.

Three days after a hindpaw injection of Complete Freund’s Adjuvant (CFA), aged mice displayed shorter withdrawal latencies than young mice in response to radiant heat (Fig. 7a). In experiments where BIBP 3226 was used to block Y1 receptors, the withdrawal latency was significantly reduced in young but not aged mice (Fig. 7b). By contrast, Y1 activation increased latencies in both young and aged mice (Fig. 7c), indicating that the antinociceptive effects of Y1 signalling are lost during aging following a reduction in the availability of endogenous NPY rather than reduced expression of functional Y1 receptors.

**Figure 7:**
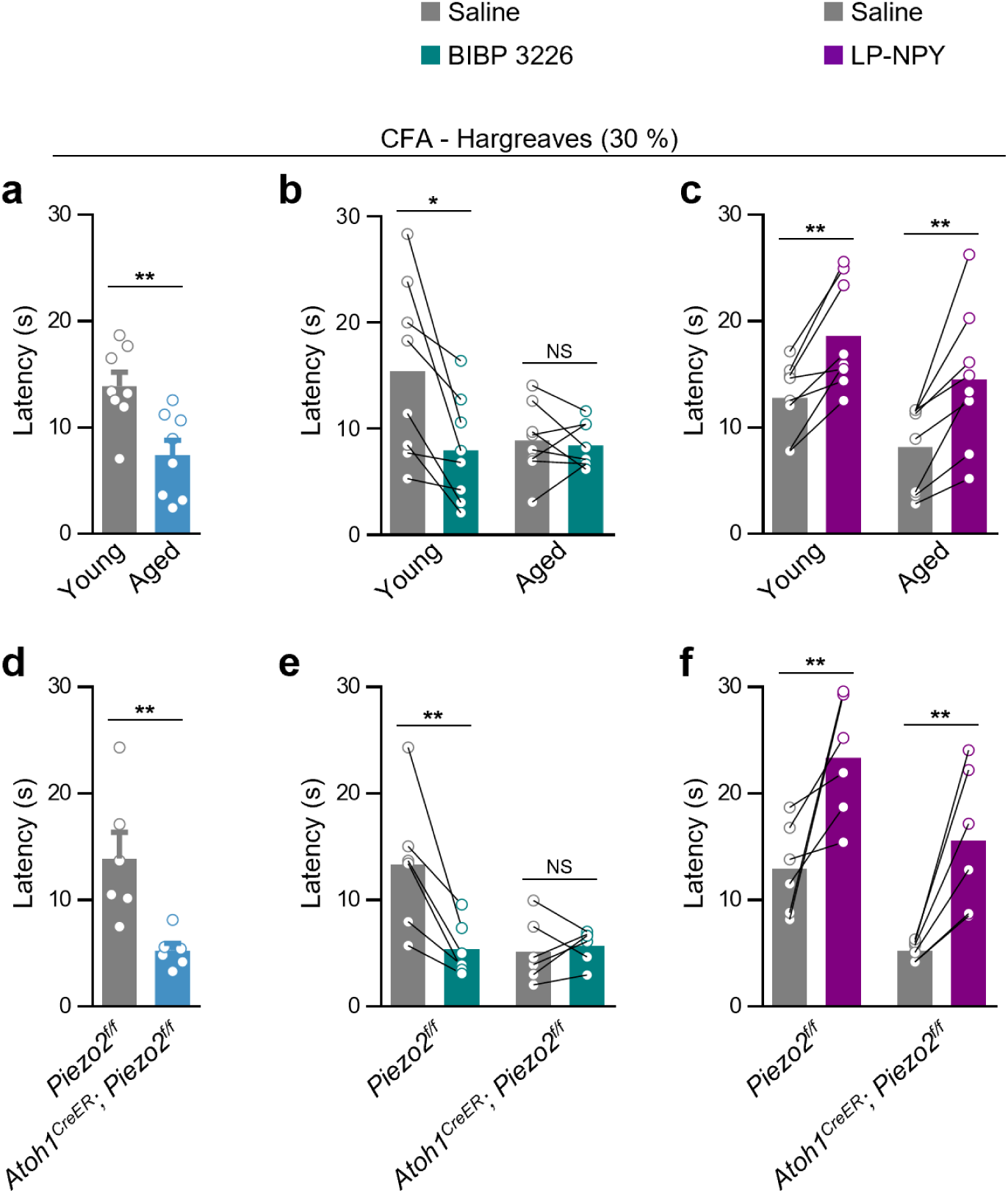
NPY^+^ neurons gate thermal allodynia in young mice but not following the loss of Merkel cell signalling. **a,** Aged mice display more severe radiant heat hyperalgesia than young mice following induction of chronic inflammatory pain by injection of CFA into the plantar hindpaw. **b, c,** Following CFA treatment, radiant heat hyperalgesia is exacerbated by Y1 blockade in young but not aged mice (**b**), but alleviated by Y1 activation in both young and aged mice (**c**). **d,** *Atoh1^CreER^; Piezo2^f/f^* mice display more severe radiant heat hyperalgesia than control *Piezo2^f/f^* mice following CFA treatment. **e,** Following CFA treatment, radiant heat hyperalgesia is exacerbated by Y1 blockade in control mice but not following the chronic persistent silencing of Merkel cells. **f,** Following CFA treatment, radiant heat hyperalgesia is alleviated by Y1 activation in both control and Merkel cell-silenced mice. Error bars represent SEM. *p < 0.05; **p < 0.01; ns, no significant difference.

We then asked whether the increased sensitivity to radiant heat seen in aged mice is caused by the loss of Merkel cell input by assessing pain responses in CFA-treated mice following the chronic silencing of Merkel cells. In CFA-treated *Atoh1^CreER^; Piezo2^f/f^* mice, the latency of responses to radiant heat was significantly reduced (Fig. 7d). By contrast, acute chemogenetic silencing of Merkel cells in CFA-treated *Atoh1^CreER^; Cdx2::FlpO; R26^ds-hM4D^* mice failed to alter the withdrawal latency to radiant heat (Extended Data Fig. 5a). These data suggest that, as with mechanical itch, Merkel cell signalling of tactile information is not required for the gating of inflammatory thermal pain, but instead that thermal hyperalgesia may be exacerbated by a central mechanism following the persistent loss of input from Merkel cells.

Lastly, we assessed whether pain responses are altered by blocking NPY-Y1 receptor signalling. Following Y1 blockade, withdrawal latency was significantly reduced in tamoxifen-treated *Piezo2^f/f^* control mice, but not in *Atoh1^CreER^; Piezo2^f/f^* mice, demonstrating that pain signalling by endogenous NPY is reduced when Merkel cells are chronically silenced (Fig. 7e). By contrast, withdrawal latencies were increased by pharmacological activation of Y1 receptors in both groups (Fig. 7f). These data argue that the loss of NPY^+^ INs caused by the degeneration of Merkel cells during aging also results in reduced NPY-Y1 gating of inflammatory pain. Moreover, these findings exemplify a mechanism by which central circuits for the processing of somatosensory information are restructured during aging.

### Itch and pain transmission neurons are resilient to reduced input from Merkel cells during aging

Our finding that the NPY agonist LP-NPY reduces itch and pain behaviours in aged mice implies the sustained expression of functional Y1 receptors and that the glutamatergic neurons that express Y1 remain intact when Merkel cell input is lost during aging. To confirm this, we analysed Y1-expressing neurons in the dorsal horn of aged compared to young mice and in young adult mice following the silencing of Merkel cells. In both groups, the number of Y1 immunoreactive neurons (Fig. 8, a-h) and the intensity of Y1 immunofluorescence within the dorsal horn were unchanged, with similar numbers of neurons in the dorsal horn expressing *Npy1r (Y1)* mRNA (Extended Data Fig. 6, a-h). Moreover, very few Y1^+^ cells exhibited TUNEL reactivity in either aged mice or following Merkel cell silencing (Fig. 8, a-h). There was also in aged and Merkel cell-silenced mice no reduction in the number of Calcrl^+^/Gpr83^+^ neurons that relay mechanical itch information from the spinal cord to the PBN^24^ (Fig. 8, a-h; Extended Data Fig. 6, i-n). Furthermore, there was no change in the number of the Tacr1^+^ neurons that transmit acute chemical itch information^24,51^ and mediate thermal hyperalgesia^52^ or the *Grpr*-expressing neurons (Extended Data Fig. 7, a-d) that overlap with the Tacr1^+^ population and relay chemical and chronic itch information. Calretinin^+^ neurons that process chemical itch and mechanical allodynia in inflammatory pain^53–55^ were also largely unchanged in aged and Merkel cell-silenced mice. Together, these data show that the excitatory neurons that transmit pain and itch in the spinal cord are largely spared following a reduction in input to the dorsal horn from Merkel cells.

**Figure 8:**
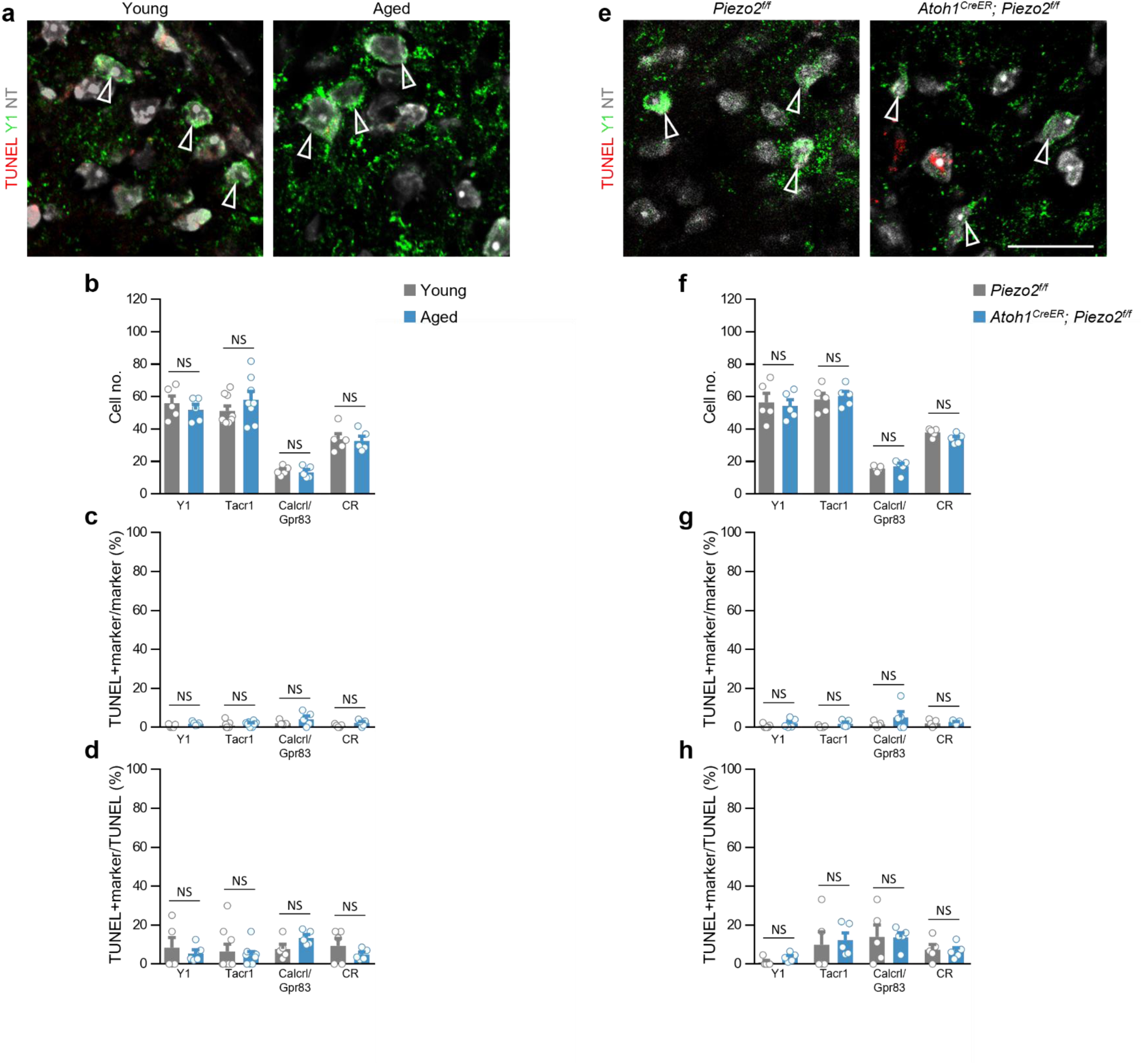
Excitatory neurons for itch and pain transmission are resilient to the loss of Merkel cell activity. **a,** Representative images showing a lack of TUNEL staining in Y1-immunoreactive neurons in the dorsal horn at cervical segment C5 of young (left) and aged mice (right). **b-d,** Programmed cell death in populations of molecularly defined excitatory neurons identified by antibodies against Y1, Tacr1, Calcrl and Gpr83, and calretinin (CR) in aged compared to young mice: summaries of population size (**b**), the proportion of TUNEL^+^ neurons expressing each molecular marker (**c**), and the proportion of each molecularly defined population labelled by TUNEL (**d**). **e,** Representative images showing a lack of TUNEL staining in Y1-immunoreactive neurons in the dorsal horn of control (left) and Merkel cell-silenced mice (right). **f-h,** Programmed cell death in populations of molecularly defined excitatory neurons identified by immunohistochemistry in Merkel cell-silenced mice compared to controls: summaries of population size (**f**), the proportion of TUNEL^+^ neurons expressing each molecular marker (**g**), and the proportion of each molecularly defined population labelled by TUNEL (**h**). Scale bars, 20 µm. Error bars represent SEM. ns, no significant difference.

## DISCUSSION

In this study we show that excitatory input from Merkel cell touch receptors is required to maintain the dorsal horn circuitry that determines mechanical itch and pain sensitivity. Specifically, our results reveal that the programmed cell death of NPY-expressing neurons is a feature of aging that is caused by the loss of input from Merkel cells. The resulting loss of NPY tone in aged animals drives the dysregulation of itch transmission by sensitizing mice to mechanical itch stimuli and exacerbating chronic itch. The antinociceptive effects of NPY-Y1 signalling are also lost following the activity-dependent degeneration of NPY^+^ INs, and this results in thermal hyperalgesia during inflammation.

### Activity-dependent regulation of neuron survival in the adult spinal cord

Our data reveal a critical homeostatic role for touch signalling by Merkel cells in maintaining the cellular composition and functional integrity of the somatosensory circuits that gate itch and pain. This role mirrors in adults the early developmental contribution that active touch makes to LTMR end organ morphogenesis^56^ and the functional wiring-up of sensorimotor reflex circuits in the spinal cord^7,8^. The activity-dependent control of neuronal cell survival has been observed in other neuronal populations during development^57–59^. It has also been proposed to occur in the adult CNS^60–62^; however, in these studies it is possible that excitotoxicity or other mechanisms contributed to cell death. The genetic approach used here selectively avoids the excitotoxic glutamate release that occurs following surgical deafferentation^61,63,64^ and instead reveals an activity-dependent mechanism that is likely mediated by Merkel cell-SAI afferents that directly innervate dorsal horn neurons in the LTMR-RZ. This conclusion is supported by our demonstration that the NPY^+^ IN population receives direct innervation from Merkel cell afferents (Extended Data Fig 4; Ref. 23)

### Central somatosensory circuits are reconfigured during aging

Touch sensitivity and protective itch- and pain-related behaviours are dynamically regulated in response to injury and disease, underscoring a need for homeostatic mechanisms that ensure the appropriate equilibration of touch-driven protective behaviours over the lifetime of the animal. We propose that impairment of activity in the pathways that transmit innocuous touch can be to some extent compensated for by heightened activity in the pathways that mediate mechanical itch and pain, which are quiescent in healthy animals. In this way, protective and corrective behaviours driven by innocuous touch can to some extent be rescued. The use of neural activity to adjust the cellular composition of somatosensory circuits in the dorsal horn is particularly well suited to this role. However, this mechanism carries the cost of aversive sensation, and the loss of cutaneous LTMR input in older animals renders permanent changes in the central circuits that transmit and gate touch, itch and pain. In support of this model, reductions in tactile acuity related to aging^18,65–69^ are associated with the degeneration of mechanosensory end organs and sensory neurons^15,65,70–72^ and with increased sensitivity to pain and itch stimuli^14,17,18,20,21^. Further support for the activity-dependent modulation of touch sensitivity and protective reflexes comes from our results showing the loss of neurons in the dorsal horn of aged mice is largely reproduced by experimental silencing of Merkel cells in young adult animals (Fig. 2). Moreover, the observation that the mechanical sensitivity of nociceptive C-fibres is not elevated in aged mice that exhibit behavioural sensitization to mechanical stimuli following inflammatory insult^73^ provides further evidence for changes to the central processing of touch information during aging, and it agrees with our finding that the central gating of inflammatory pain is impaired in aged mice.

### Differential loss of defined populations of dorsal horn neurons during aging

In both Merkel cell-silenced and aged mice, the loss of neurons by apoptosis is largely restricted to the LTMR-RZ, where SAI fibres terminate, whereas lamina I doesn’t receive direct SAI input and was completely spared (Fig. 2). Even though reductions were seen in the number of excitatory neurons, we did not observe cell loss in neuronal cell types that are known to transmit mechanical and chemical itch, thereby indicating reduced inhibition as the likely driver of increased itch in aged mice. Tellingly, the NPY^+^ INs that gate mechanical itch as opposed to the dynorphin^+^ neurons that gate chemical itch are reduced in number following the loss of Merkel cell signalling (Fig. 3). Although these populations have overlapping laminar distributions, the NPY^+^ INs are highly concentrated in the LTMR-RZ, while the majority of dynorphin^+^ neurons are located more superficially in laminae that receive little in the way of input from Merkel cell-SAI afferent complexes. Moreover, the NPY^+^ INs receive direct input from SAI afferents, supporting our conclusion that the survival of NPY^+^ INs is dependent on monosynaptic input from Merkel cells-SAI complexes.

Our observation that other as-yet unidentified subsets of excitatory neurons and NPY-negative inhibitory interneurons also degenerate following the loss of Merkel cell input suggests that age-related activity-dependent cell loss in the dorsal horn may contribute to other somatosensory deficits that are common in aging^65–69,74–76^. Furthermore, our finding that excitatory input to the dorsal horn from touch receptors supports the survival of molecularly defined subsets of neurons in the dorsal horn has implications not only for somatosensory dysfunction during normal aging, but also for somatosensory dysfunction associated with the peripheral neuropathies that occur in diseases such as diabetes mellitus or Charcot-Marie Tooth Disease, or as a side effect of chemotherapy.

### NPY-Y1 neuropeptide signalling in itch and pain

Here we have shown that the loss of spinal NPY^+^ neurons and NPY inhibitory tone that occurs during aging alters activity in at least two separate somatosensory transmission pathways within the spinal cord, both of which are modulated by NPY-Y1 signalling^22–26,40,52,77^. These changes underlie at least two forms of sensory dysfunction associated with old age, namely mechanical hyperknesis/chronic itch and thermal hyperalgesia that is associated with inflammatory pain^17,20^.

Mechanical itch is gated predominantly by a peptidergic mechanism rather than by fast inhibitory transmission, and NPY-Y1 signalling is hypothesized to determine the sensitivity of the mechanical itch pathway to touch stimuli^22^. Our finding that chemogenetic silencing of NPY^+^ INs potentiates mechanical itch responses, but not in the presence of a selective Y1 inhibitor (Fig. 5)^22^, agrees with our previous report that pharmacological blockade of Y1 receptors or selective deletion of *Npy1r* from dorsal horn neurons potentiates mechanical itch^22^. Likewise, chronic inflammatory pain is modulated by endogenous NPY-Y1 signalling in young adult rodents, such that knockout or blockade of Y1 receptors exacerbates mechanical and thermal hyperalgesia following inflammation in young mice^25,26,78,79^. The loss of NPY tone that is due to the loss of Merkel cells during aging accounts for the chronic itch phenotype observed in geriatric mice and may therefore account for the increased susceptibility of older humans to chronic itch conditions^20^. Chronic itch is sustained by a vicious cycle of itching, scratching and inflammation^80^ that is exacerbated by hypersensitivity to mechanical stimuli (hyperknesis/alloknesis)^22,23^.This accords with our previous demonstration that dermatological chronic itch is driven by simultaneous activation of the NPY-sensitive pathway that conveys mechanical itch and an NPY-insensitive neural pathway that normally conveys chemical itch signals mechanical itch^24^. Similarly, disinhibition of pain transmission pathways following the loss of NPY-Y1 signalling likely contributes to the increased prevalence of chronic pain conditions and the sensitization to painful stimuli that occurs during aging^17^. Strikingly, the loss of NPY^+^ INs that occurs following Merkel cell silencing or the loss of Merkel cells that occurs during normal aging is partial, phenocopying the 60-65% reduction in the spinal NPY^+^ IN population induced by intersectional diphtheria toxin receptor-mediated ablation, which also causes a severe chronic itch phenotype characterized by hyperknesis and spontaneous scratching^22–24,40^.

Finally, while our findings show that these itch and pain pathways become insensitive to endogenous NPY following the loss of NPY+ INs during aging, the Y1 receptor remains a potential therapeutic target for chronic itch and pain in older patients, since the Y1-expressing glutamatergic neurons that form these pathways are unchanged in number, and pharmacological activation of Y1 receptors suppresses itch and pain in aged mice (Fig. 5, 8). Thus, understanding how the architecture of the dorsal horn changes throughout the lifespan may prove to be indispensable when designing pharmacological strategies for the treatment of somatosensory disorders in older patients.

## Methods

### Tissue Preparation for Histology

Mice were euthanized by a single intraperitoneal (i.p.) injection (10 µl g^-1^ body weight) of ketamine (10 mg ml^-1^) and xylazine (1 mg ml^-1^) immediately prior to perfusion with 20 ml ice-cold 4% paraformaldehyde (PFA) in PBS. Spinal cords were dissected and post-fixed for 1 h at RT, then washed 3 times in PBS. Skin was depilated and dissected, fixed overnight in 4% PFA at 4°C, then washed 3 times in PBS. All tissues were then cryoprotected in 30% sucrose-PBS (w/v) overnight at 4 °C before being embedded in Tissue-Tek OCT Compound (Sakura Finetek) and cryosectioned at 14 µm or 30 µm. For histological analysis, sections were dried at RT and stored at -20 °C.

### TUNEL Labelling

*In situ* cell death detection kit TMR red (Roche 12156792910) was used to detect cell death using TUNEL technology following the manufacturer’s instructions. Briefly, spinal cord sections were washed with PBS for 30 min at room temperature and permeabilized with 0.1% Triton X-100, 0.1% sodium citrate for 2 min on ice. Sections were washed twice with PBS and incubated with the TUNEL mix (label solution + enzyme terminal transferase) for 60 min at 37 °C in a humidified chamber. Sections were then stained with antibodies by the method below.

### Immunohistochemistry

The following primary antibodies were used in this study: rabbit α-Calcrl (1:50; Invitrogen), rabbit α-Calretinin (1:250; Swant), rabbit α-Caspase 3 (1:500; Invitrogen), rabbit α-DsRed (1:1000; Clontech), rabbit α-Galanin (1:500; Peninsula Lab), chicken α-GFP (1:000; Aves), rabbit α-Gpr83 (1:500; Alomone Labs), rat α-K8 (TROMA1, Developmental Studies Hybridoma Bank, 1:100), rabbit α-NF200 (1:1000; Sigma), mouse α-NF200 (1:500; Sigma), rabbit α-NK1R (1:500; Advanced Targeting Systems), rabbit α-NPY (1:500; Peninsula Lab), rabbit α-NPY1R (1:500; Neuromics), guinea pig α-prodynorphin (1:500; Abcam), rabbit α-Parvalbumin (1:1000; Swant), rabbit α-Pax2 (1:200; Zymed), rat α-RFP (1:1000; Chromotek), rabbit α-dsRed (1:1000 Clontech), guinea pig α-vGluT1 (1:1000; Millipore). In addition, Alexa Fluor 647-conjugated isolectin GS-IB4 from *Griffonia simplicifolia* (Invitrogen) was used at 1:500.

For sectioning, tissues were cryoprotected in 30% sucrose-PBS (w/v) overnight at 4 °C before being embedded in Tissue-Tek OCT Compound (Sakura Finetek) and cryosectioned at 14 µm or 30 µm. Sections were dried at RT and stored at -20 °C. Sections were then washed once with PBS (5 min), blocked with a solution of 10% donkey serum in PBT (PBS, 0.1% Triton X-100) for 1 h at RT and then incubated overnight at 4 °C with primary antibodies in a solution of 1% donkey serum in PBT. Sections were then washed 3 times (15 min each) in PBT before being incubated for 2 h at RT with fluorophore-conjugated secondary antibodies (1:1000; Jackson Laboratories) in a solution of 1% donkey serum in PBT. Sections were then washed 3 times (15 min each) in PBT. Some sections were then incubated with DAPI for 1 min or NeuroTrace 435/455 (1:300 in PBT; Invitrogen) for 30 min before being washed once in PBT and mounting in Aqua-Poly/Mount (Polysciences).

For whole-mount skin preparations, fixed skin from the nape and rostral back was cut into small pieces and washed with PBS containing 0.3% Triton X-100 (0.3% PBT) every 10 min for 2 hr. Then, the skin was incubated with primary antibodies in 0.3% PBT containing 5% donkey serum and 20% DMSO at RT for 5 days. Tissues were then washed with 0.3% PBST every 10 minutes for 2 hours and incubated with secondary antibodies and DAPI in 0.3% PBST containing 5% donkey serum and 20% DMSO and at RT for 3 days. Finally, tissues were washed with 0.3% PBST every 10 minutes for 2 hours before being mounted with Aqua-Poly/Mount (Polysciences).

A Zeiss LSM 700 confocal microscope was used to capture images. ImageJ software was used to assess immunofluorescence, with thresholds set according to signal intensity^81^. For quantification, three cervical sections from each spinal cord were analysed per condition. Cross sectional area of the dorsal horn did not differ between young and aged mice (young, n =4; aged, n =3; *p* > 0.05) or control *Piezo2^f/f^* and *Atoh1^CreER^; Piezo2^f/f^* mice (*Piezo2^f/^*^f^, n = 3; Atoh1^CreER^*; Piezo2^f/f^*, n = 4; *p* > 0.05).

### Rabies Virus Tracing

For the transsynaptic tracing studies, P10 *NPY::Cre; Lbx1^FlpO^; R26^ds-HTB^* mice were anesthetized with isoflurane and placed on a stereotaxic frame. A skin incision in the back was made to expose cervical the spinal cord. EnvA-pseudotyped, ΔG-mCherry (250 µl, 3.3 x 1010 units ml-1) was injected starting from 300 μm deep into the dorsal horn and going back dorsally, in 6 steps consisting of 50 nl pulses every 50 μm on the left side (400 μm lateral from the midline) of the spinal cord using a 0.5 μl Hamilton syringe mounted on a UMP3 UltraMicroPump (WPI). Skin was then sutured with a nylon surgical suture. Tissue was collected at P18.

### RNA Fluorescence *In Situ* Hybridization

For RNAscope RNA FISH, the following probes were used: Mm-Slc32a1, Mm-Slc17a6-C3, Mm-Grpr-C2 (Advanced Cell Diagnostics). These were revealed with Opal 520, Opal 520, Opal 650 (PerkinElmer).

Tissue was prepared as described above for immunohistochemistry and cryosectioned at 14 µm. Multiple-labelling fluorescence *in situ* hybridization was performed using the RNAscope Multiplex Fluorescent Reagent Kit v2 Assay (Advanced Cell Diagnostics) following the manufacturer’s recommended protocol. Sections were washed with DAPI for 1 min and mounted Aqua-Poly/Mount (Polysciences). Sections were then imaged under a 40x oil objective on a Zeiss LSM700 confocal microscope. For quantification, three sections from each spinal cord were analysed per condition.

### Cell Ablation

For ablations using saporin-conjugated receptor ligands, 2-year-old mice were given a single i.t. injection of PEN-saporin (PEN-SAP; 3 µg in 10 μl 0.9% sterile saline) to ablate Gpr83^+^ neurons or [Sar^9^, Met(O_2_)^11^]-substance P-saporin (SSP-SAP; 200 ng in 10 μl 0.9% sterile saline; Advanced Targeting Systems)^22,24,82^ to ablate Tacr1^+^ neurons. PEN-SAP was produced by CytoLogistics (Carlsbad, CA) by biotinylation of PEN peptide, the endogenous ligand of Gpr83, followed by incubation with streptavidin-ZAP (Advanced Targeting Systems)^83,84^. Littermate controls received injections of an equal mass of blank saporin (Advanced Targeting Systems) in 10 μl 0.9% sterile saline. Behavioural testing was performed 14 days later.

### Drug Administration

Synthesis of the selective Y1 receptor agonist [Leu^31^, Pro^34^]-Neuropeptide Y (LP-NPY; YPSKPDNPGEDAPAEDMARYYSALRHYINLLTRPRY-NH2)^50^ was performed on a Gyros Protein Technologies, Inc. Tribute peptide synthesizer equipped with real-time UV monitoring, using standard Fmoc chemistry, in the Salk’s Peptide Synthesis Core^22^. The resultant crude peptide was purified by the Salk’s Proteomics Core to > 98% using HPLC. The final, purified product gave a single peak of predicted mass (4240.7 Da) by MS analysis. The peptide was administered in solution at pH 7.

For i.t. injections, mice were briefly anesthetized with 2.5% isoflurane in O_2_, and a 30-gauge needle was inserted into the fifth intervertebral space until it elicited a tail flick. The needle was held in place for 30 s and turned 90° prior to withdrawal to prevent outflow.

[Leu^31^, Pro^34^]-NPY was dissolved in 0.9% sterile saline. Clozapine *N*-oxide (CNO; Sigma) and the Y1 receptor antagonist BIBP 3226 (Tocris), were dissolved in DMSO, which was then diluted with so that the concentration of DMSO did not exceed 1% in injected solutions.

To activate Cre, Cre^+^ mice and Cre^-^ controls received an i.p. injection of tamoxifen at 100 mg kg^-1^ body weight once per day for 5 consecutive days from P56 ^14^ by which time stem cells are quiescent. Experiments were performed between 7 and 14 days after tamoxifen injection.

CNO was administered by i.p. injection at 2 mg kg^-1^. BIBP 3226 (0.2 ng kg^-1^) and LP-NPY (1 ng kg^-1^) were administered by i.t. injection. Mice were tested 15-45 min following administration.

### Inflammatory Pain Induction

To induce inflammatory pain, mice were briefly anesthetized with isoflurane (3–5 min at 2%), and 20 μl Complete Freund’s Adjuvant (CFA, Sigma) was injected into the plantar surface of the left hindpaw. Responses to radiant heat (30% beam intensity) were assessed 3 days after CFA treatment.

### Behavioural Testing

Littermate controls were used for behavioural tests, and the experimenter was blinded to genotype/treatment. Animals were habituated to the behavioural testing apparatus for 1 h on each of the two days preceding data collection, and for 30 min on the day of testing.

#### Spontaneous Itch

To quantify scratching induced in the absence of a mechanical stimulus (spontaneous itch), mice were placed in a plastic chamber and video-recorded for a period of 30 min. Bouts of hindlimb scratching were counted offline^22,23^.

#### Mechanical Itch

To quantify itch-related scratching behaviours induced by mechanical stimulation of the hairy skin, mice were placed in a plastic chamber and each of a series of low-grade Frey hairs was applied to the nape for 1 s at an interval of 1 min. Mice were stimulated 5 times with each filament, and the number of trials in which scratching was evoked was recorded^22,23,48^.

#### Hargreaves

On day 3 after CFA treatment to induce inflammatory pain, mice were placed in a plastic chamber and the plantar surface of the left hindpaw was exposed to a beam of radiant heat (30% maximum intensity; IITC, USA). The latency to paw withdrawal was determined in over 3 trials and averaged per animal, with a 5 min interval between trials. A cutoff time of 30 s was set to prevent tissue damage.

#### von Frey

Mice were placed in a plexiglass chamber on an elevated wire grid and the lateral plantar surface of the hindpaw was stimulated with calibrated von Frey monofilaments (0.008-4 g). The paw withdrawal threshold for the von Frey assay was determined by Dixon’s up-down method^85^.

#### Brush

Mice were placed in a plexiglass chamber on an elevated wire grid and the plantar surface of the hindpaw was stimulated by light stroking with a fine paintbrush in a heel-to-toe direction^44^.The test was repeated 10 times at 10 s intervals between trails, and the percentage of positive paw-withdrawal trials was calculated.

#### Pinprick

Mice were placed in a plastic chamber on an elevated wire grid and the plantar surface of the hindpaw was stimulated with an Austerlitz insect pin (Tip diameter: 0.02 mm; Fine Science Tools). The pin was gently applied to the plantar surface of the hindpaw without moving the paw or penetrating the skin. The pin stimulation was repeated 10 times on different paw areas with a 1-2 min interval between trails, and the percentage of trials in which mice responded with paw withdrawal was calculated.

### Alloknesis and chronic itch models

#### Histamine alloknesis

Histamine (50 µg in 10 µl) was administered by intradermal injection into the nape 30 min prior to testing, as previously described^48^. A 0.008 g or 0.07 g von Frey hair (as indicated) was then used to deliver light punctate stimuli to a randomly selected site 5-10 mm from the injection site. Mice were assessed for hindlimb scratching immediately following each stimulus. Mice received 10 stimulations at 1 min intervals, and data were presented as the number of trials in which scratching was evoked. BIBP 3226, LP-NPY or control saline were administered by i.t. injection immediately prior to histamine injection.

#### Dry skin itch model

The nape and rostral back of P56 or 2-year-old mice were shaved 5 days before treatment. Treatment entailed application with a sterile cotton swab of a mixture of acetone and ether (AEW) in a ratio of 1:1 to the rostral back for 15 s, followed by distilled water for 25 s^46,47^. AEW treatment was repeated twice daily on days 1-7. On day 8, mechanical itch sensitization was assessed. Mice were first habituated for 15 min in a plexiglass chamber, and then a 0.07g von Frey filament was used to deliver 10 innocuous mechanical stimuli (∼1-s duration) at an interval of 1 min to a randomly selected site on the nape at the margin of the AEW-treated area. The response rate of hindlimb scratching toward the stimulation site was quantified as a percentage. Mice continued to receive AEW treatment twice daily from day 8 to day 10. On day 10, spontaneous itch was measured as described above.

### Quantification and Statistical Analysis

Data were analysed in GraphPad Prism 9.2 (Graphpad Software) or Excel 16 (Microsoft) by two-way ANOVA with Šidák post hoc correction, or by two-tailed t-tests. *p* < 0.05 was considered to be statistically significant. All data are presented as the mean ± standard error of the mean (SEM).

## Acknowledgments

This study was supported by NIH grant NS111643 to M.G. Viruses were supplied by the Salk Institute Viral Core Facility. M.G. holds the Frederick W. and Joanna. J. Mitchell Chair.

## Author contributions

DA and MG designed the experiments. DA and SP performed the experiments. DA and MG wrote and edited the manuscript.

