## Extended Data for "Age-related sensory dysfunction reconfigures the spinal circuitry for touch, itch and pain"

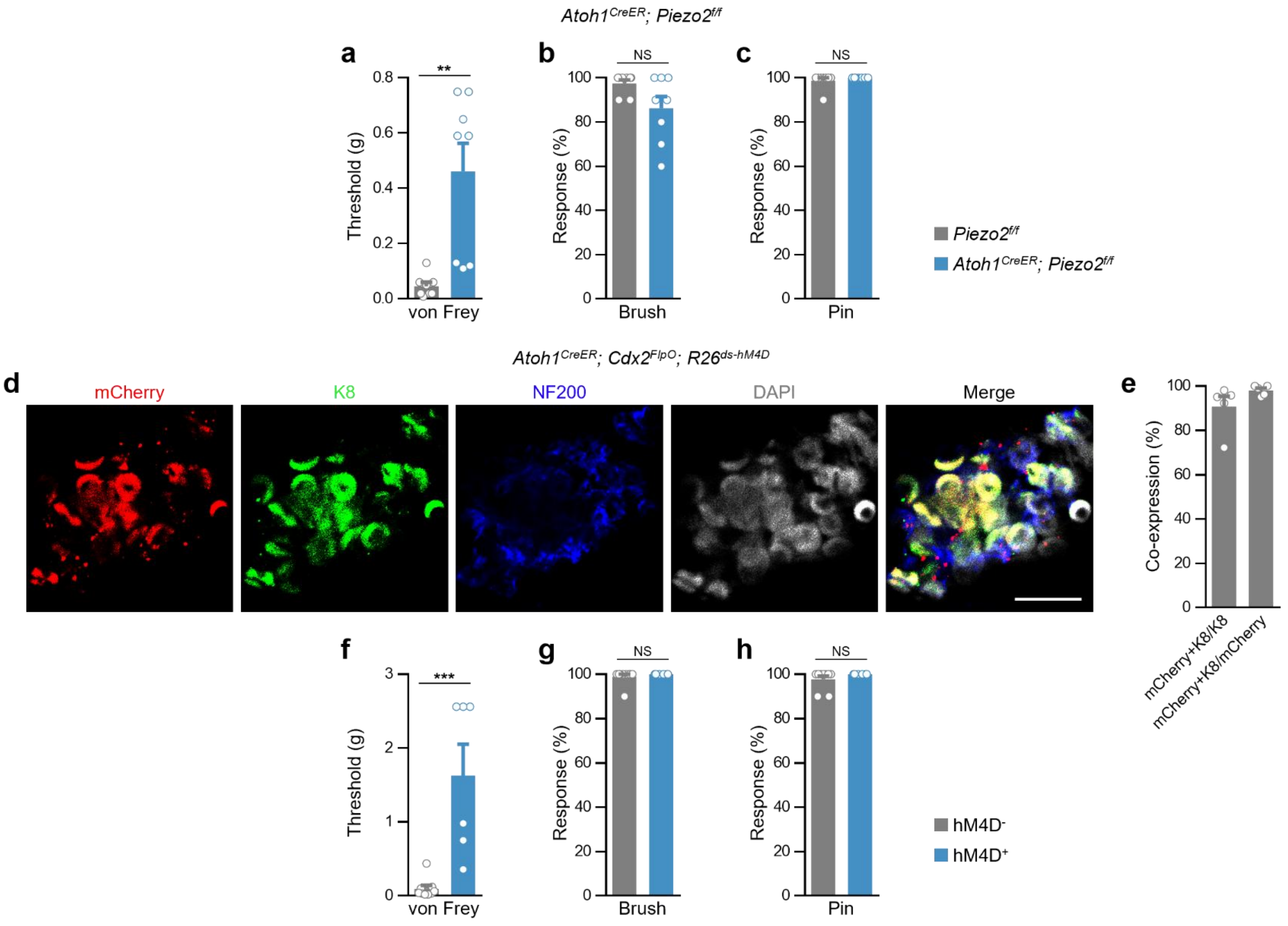
Atoh1^CreER; Piezo2^f/f
a
b
c
**
NS
NS
Threshold (g)
Response (%)
von Frey
Brush
Pin
Piezo2^f/f
Atoh1^CreER; Piezo2^f/f
Atoh1^CreER; Cdx2^FlpO; R26^ds-hM4D
d
mCherry
K8
NF200
DAPI
Merge
e
Co-expression (%)
mCherry+K8/K8
mCherry+K8/mCherry
f
g
h
***
NS
NS
Threshold (g)
Response (%)
von Frey
Brush
Pin
hM4D-
hM4D+

**Extended Data Figure 1: Chronic but not acute loss of Merkel cell function exacerbates itch.**

**a-c,** Chronic silencing of Merkel cells in *Atoh1*$^{CreER}$*; Piezo2*$^{f/f}$ mice reduces sensitivity to static touch (**a**) but not dynamic touch (**b**) or acute pain (**c**). **d, e,** Representative images of a whole-mount hairy skin preparation from *Atoh1*$^{CreER}$*; Cdx2*$^{FlpO}$*; R26*$^{ds\text{-}hM4D}$ mice showing mCherry expression in antibody-labelled $K8^+$ Merkel cells (**d**) and summary (**e**) of mCherry and K8 co-expression. Scale bar, 10 µm. **f-h,** Acute silencing of Merkel cells in *Atoh1*$^{CreER}$*; Cdx2*$^{FlpO}$*; R26*$^{ds\text{-}hM4D}$ mice compared to *Atoh1*$^{CreER}$*; R26*$^{ds\text{-}hM4D}$ controls following CNO administration reduces sensitivity to static touch (**f**) but not dynamic touch (**g**) or acute pain (**h**). Error bars represent SEM. $^{**}p < 0.01$; $^{***}p < 0.001$; ns, no significant difference.

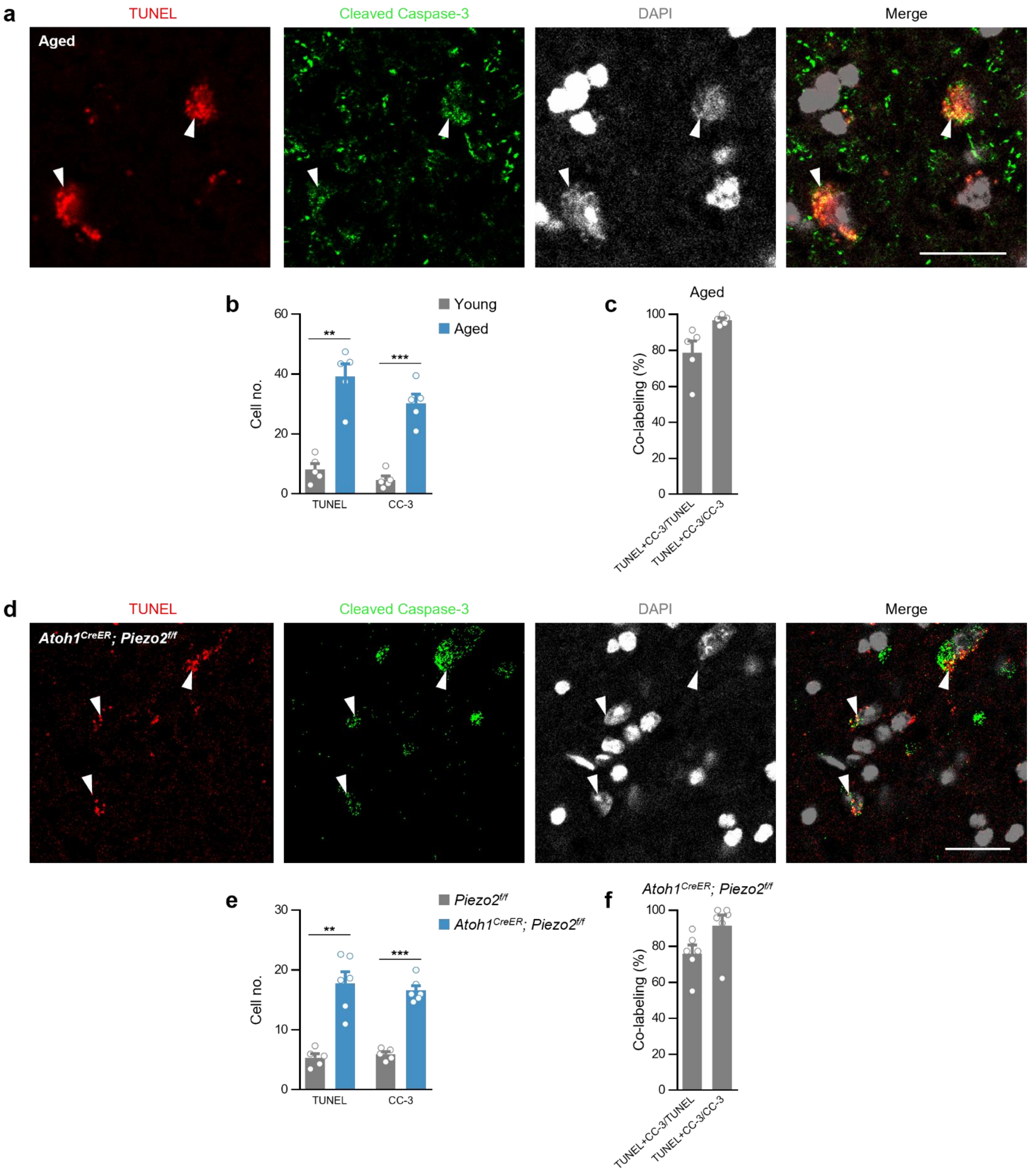

a
TUNEL
Cleaved Caspase-3
DAPI
Merge
Aged
b
Young
Aged
Cell no.
60
40
20
0
**
***
TUNEL
CC-3
c
Aged
Co-labeling (%)
100
80
60
40
20
0
TUNEL+CC-3/TUNEL
TUNEL+CC-3/CC-3
d
TUNEL
Cleaved Caspase-3
DAPI
Merge
Atoh1CreER; Piezo2f/f
e
Piezo2f/f
Atoh1CreER; Piezo2f/f
Cell no.
30
20
10
0
**
***
TUNEL
CC-3
f
Atoh1CreER; Piezo2f/f
Co-labeling (%)
100
80
60
40
20
0
TUNEL+CC-3/TUNEL
TUNEL+CC-3/CC-3

**Extended Data Figure 2: Aging and Merkel cell-silencing cause apoptosis in the dorsal horn.**

**a,** Representative images from the dorsal horn of aged mice showing co-staining of nuclei with TUNEL and anti-cleaved caspase-3 (CC-3). **b,** Number of cells per section exhibiting TUNEL$^-$ and CC-3-labelling in aged compared to young mice. **c,** Summary of co-labelling by TUNEL and anti-CC-3 in aged mice. Scale bars, 20 μm. **d,** Representative images from the dorsal horn of Merkel cell-silenced mice showing co-staining of nuclei with TUNEL and anti-CC-3. **e,** Number of cells per section exhibiting TUNEL and CC-3-labelling in Merkel cell-silenced compared to control mice. **f,** Summary of TUNEL and anti-CC-3 co-labelling in Merkel cell silenced mice. Error bars represent SEM. $^{**}p < 0.01$; $^{***}p < 0.001$

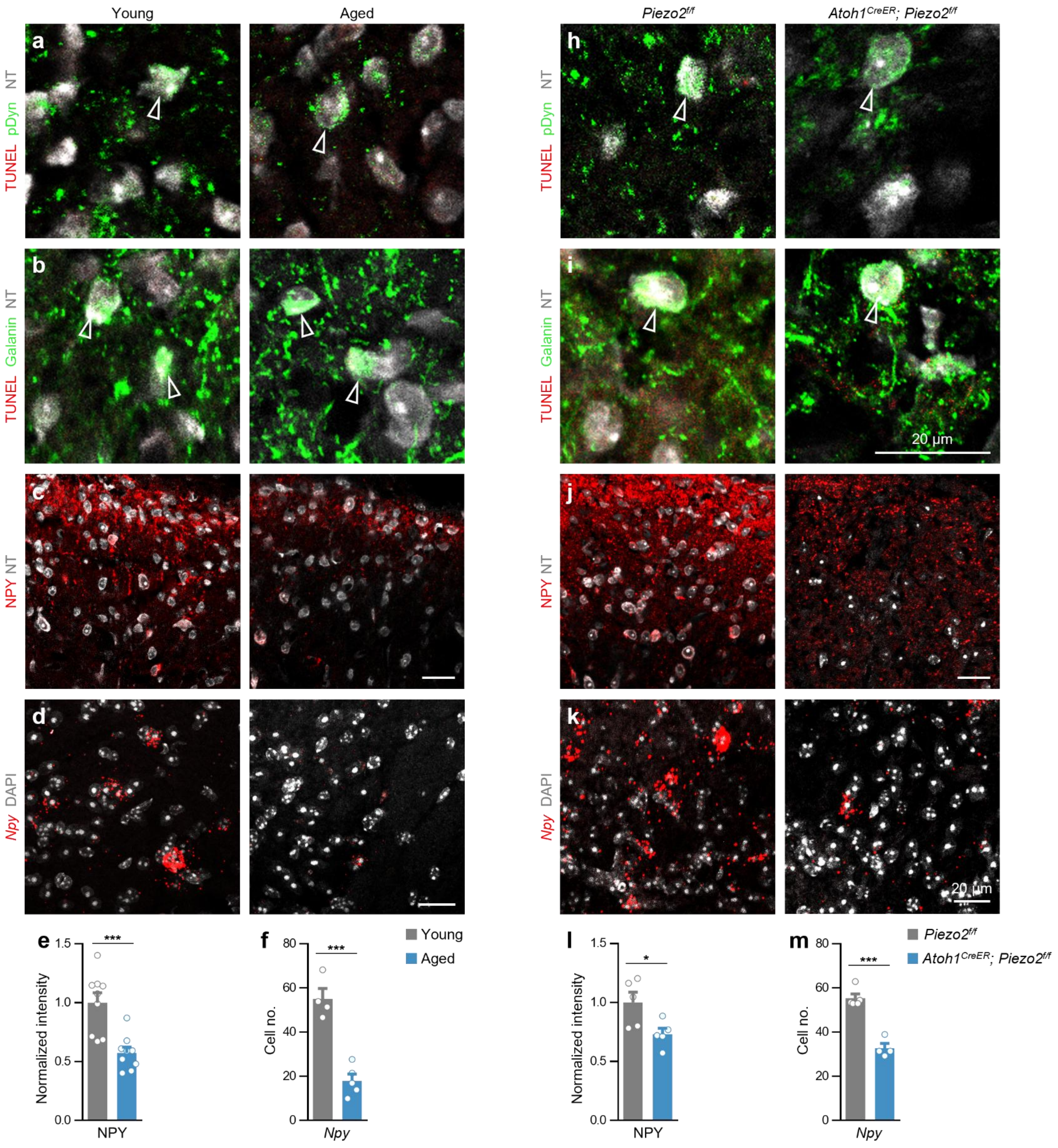

Young
Aged
Piezo2f/f
Atoh1CreER; Piezo2f/f
a
TUNEL pDyn NT
b
TUNEL Galanin NT
c
NPY NT
d
Npy DAPI
h
TUNEL pDyn NT
i
TUNEL Galanin NT
20 µm
j
NPY NT
k
Npy DAPI
20 µm
e
Normalized intensity
NPY
***
f
Cell no.
Npy
***
Young
Aged
l
Normalized intensity
NPY
*
m
Cell no.
Npy
***
Piezo2f/f
Atoh1CreER; Piezo2f/f

**Extended Data Figure 3: Reduced NPY expression following the chronic loss of Merkel cell activity.**

**a, b,** Representative images showing a lack of TUNEL labelling (open arrowheads) in dynorphin (pDyn)$^{+}$ (**a**) and galanin$^{+}$ neurons (**b**) in the dorsal horn at cervical segment C5 of aged (right) compared to young mice (left). **c, d,** Sections through the dorsal horn showing reduced NPY immunoreactivity (**c**) and *Npy* mRNA expression (**d**) in aged mice. **e,** Normalized intensity of NPY immunofluorescence is reduced in aged compared to young mice. **f,** Expression of *Npy* mRNA is reduced in aged mice. **h, i,** Representative images showing a lack of TUNEL labelling in dynorphin$^{+}$ (**h**) and galanin$^{+}$ neurons (**i**) in the dorsal horn of *Atoh1$^{CreER}$; Piezo2$^{f/f}$* (right) compared to control *Piezo2$^{f/f}$* mice (left). **j, k,** Representative images showing reduced NPY immunoreactivity (**j**) and *Npy* expression (**k**) in the cervical dorsal horn of Merkel cell-silenced mice. **l,** NPY immunofluorescence is reduced in the dorsal horn of *Atoh1$^{CreER}$; Piezo2$^{f/f}$* mice. **m,** Expression of *Npy* mRNA is reduced in Merkel cell-silenced mice. NT, NeuroTrace. Scale bars, 20 µm. Error bars represent SEM. *$p < 0.05$; **$p < 0.01$; ***$p < 0.001$; ns, no significant difference.

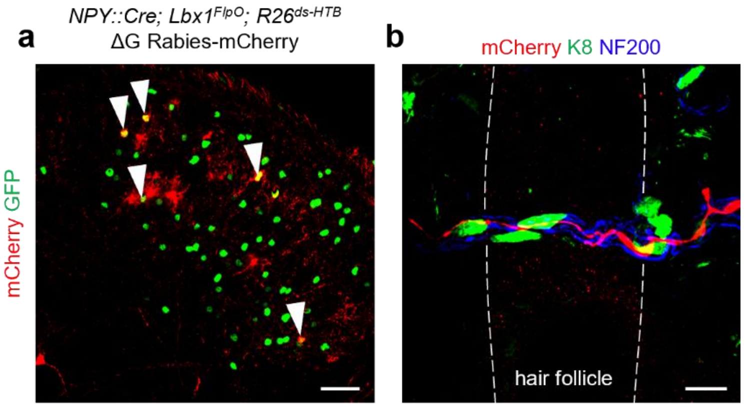


**Extended Data Figure 4: NPY⁺ INs receive input from Merkel cells in the hairy skin.**

**a,** Section through the cervical dorsal horn of a P18 *NPY::Cre; Lbx1$^{FlpO}$; R26$^{ds\text{-}HTB}$* mouse injected with EnvA G-deleted rabies-mCherry virus. Arrowheads indicate infected *NPY::Cre*$^{+}$ neurons. mCherry$^{+}$/GFP$^{-}$ cells represent transsynaptically labelled presynaptic neurons. Scale bar, 50 μm. **b,** Section of hairy skin stained with antibodies against mCherry (red), K8 (green), NF200 (blue). Scale bar, 10 μm.

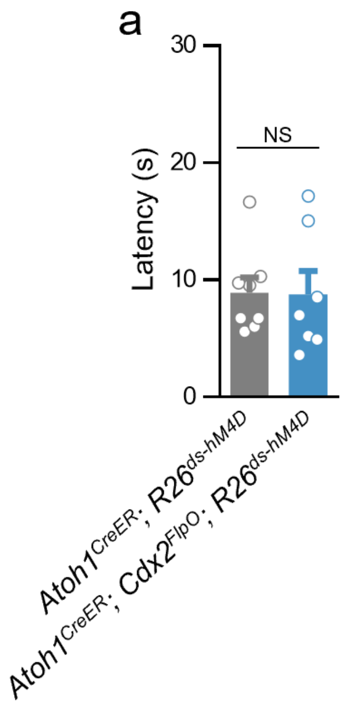


**Extended Data Figure 5: Acute silencing of Merkel cells does not alter thermal hyperalgesia.**

**a,** Response latencies to radiant heat are unaltered in CFA-treated *Atoh1*$^{CreER}$*; Cdx2*$^{FlpO}$*; R26*$^{ds\text{-}hM4D}$ mice compared to *Atoh1*$^{CreER}$*; R26*$^{ds\text{-}hM4D}$ controls following CNO administration to silence Merkel cells acutely. Error bars represent SEM. ns, no significant difference.

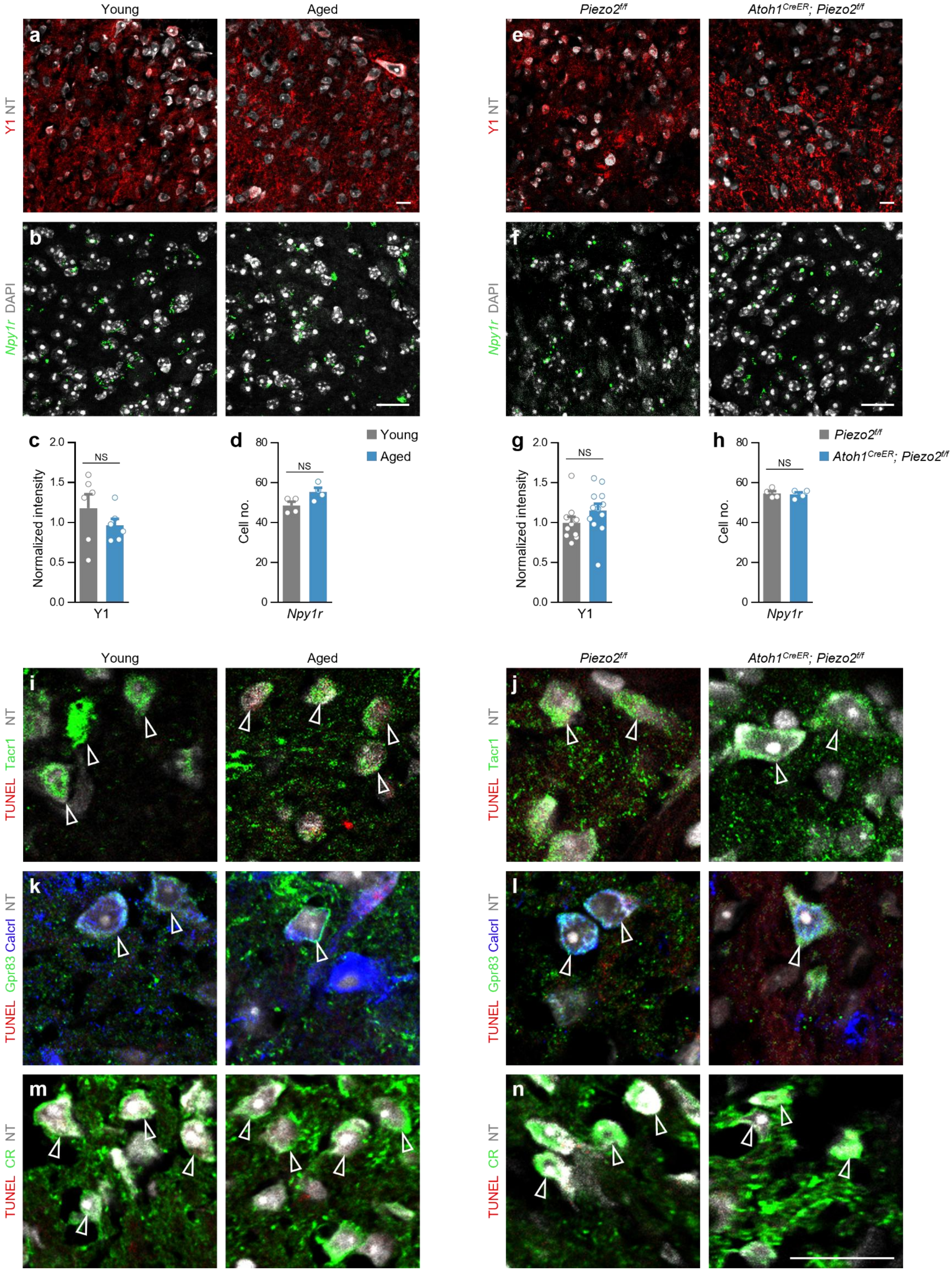

Young
Aged
Piezo2f/f
Atoh1CreER; Piezo2f/f
a
Y1 NT
b
Npy1r DAPI
e
f
c
Normalized intensity
Y1
NS
d
Cell no.
Npy1r
Young
Aged
g
h
Piezo2f/f
Atoh1CreER; Piezo2f/f
i
TUNEL Tacr1 NT
j
k
TUNEL Gpr83 Calcrl NT
l
m
TUNEL CR NT
n

**Extended Data Figure 6: Y1 expression does not change following loss of input from Merkel cells.**

**a, b,** Sections through the dorsal horn at cervical segment C5 of young (left) and aged mice (right) showing unchanged Y1 immunoreactivity (**a**) and expression of *Npy1r* mRNA (**b**). **c, d,** Normalized intensity of Y1 immunofluorescence (**c**) and expression of *Npy1r* mRNA (**d**) are unchanged in aged compared to young mice. **e, f,** Cervical dorsal horn sections from control (left) and Merkel cell-silenced mice (right) showing unchanged Y1 immunoreactivity (**e**) and *Npy1r* mRNA expression (**f**). **g, h,** Y1 immunofluorescence (**g**) and expression of *Npy1r* mRNA (**h**) are unchanged in the dorsal horn of $Atoh1^{CreER}$; $Piezo2^{f/f}$ mice. **i-n,** Representative images showing a lack of TUNEL labelling in $Tacr1^+$, $Gpr83^+$/$Calcrl^+$ and calretinin $(CR)^+$ neurons in the dorsal horn in aged compared to young mice (**i, k, m**) and $Atoh1^{CreER}$; $Piezo2^{f/f}$ compared to control $Piezo2^{f/f}$ mice (**j, l, n**). NT, NeuroTrace. Scale bars, 20 μm. Error bars represent SEM. ns, no significant difference.

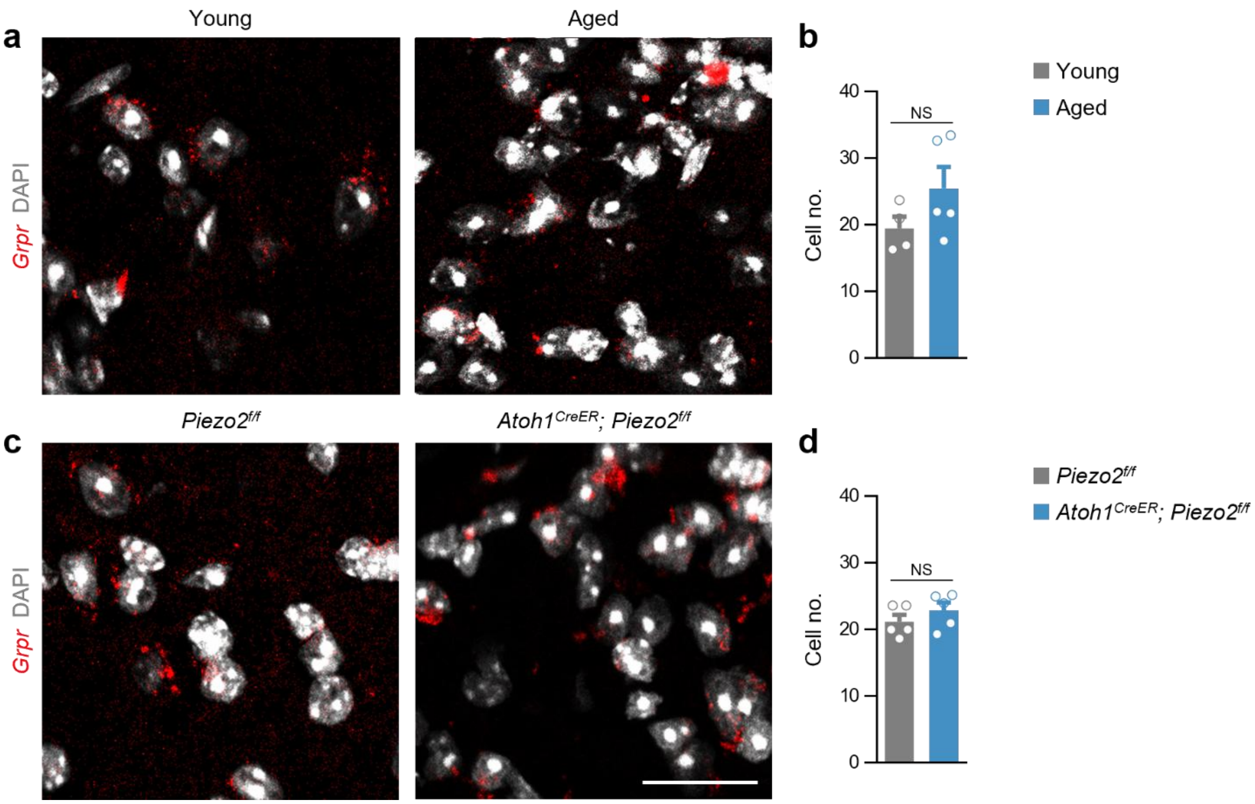


**Extended Data Figure 7: *Grpr*+ neurons are resistant to apoptosis following loss of input to the dorsal horn from Merkel cells.**

**a, b,** Representative images (**a**) and summary (**b**) showing no difference in the number of neurons expressing *Grpr*, identified by RNAScope, between young and aged mice. **c, d,** Representative images (**c**) and summary (**d**) showing no difference in the number of neurons expressing *Grpr* between *Piezo2*$^{f/f}$ and *Atoh1*$^{CreER}$; *Piezo2*$^{f/f}$ mice following tamoxifen treatment.. Scale bars, 20 µm. Error bars represent SEM. ns, no significant difference.
